# Argonaute-2 slicing of a domesticated retrotransposon maintains proteostasis and prevents lethal skeletal muscle defects

**DOI:** 10.64898/2026.05.22.727060

**Authors:** Nickolas Almodovar, Caroline Dunn, Benjamin Kleaveland

## Abstract

Argonaute (AGO) proteins associate with small guide RNAs to bind and repress mRNA targets. AGO2 is the primary AGO slicer in mammals, cleaving RNAs that base-pair extensively to its guide RNA. Disrupting the slicing activity of AGO2 in mice causes neonatal lethality, however, the specific cell types and substrates that contribute to this lethality are not known. Through a combination of genetic, histologic, and molecular approaches, we identify retrotransposon-like-1 (*Rtl1*) as a key AGO2 slicing substrate in placental endothelium and skeletal muscle, and show that AGO2 slicing in skeletal muscle, but not endothelium, is required for postnatal viability. Loss of AGO2 slicing causes cell-autonomous, pathologic changes in skeletal muscle, characterized by larger fibers, increased central nuclei, prominent central clearings, and induction of transcripts associated with heat shock proteins. Induced expression of RTL1, a domesticated retrotransposon-derived protein that can self-assemble into viral-like capsids, activates a similar heat shock protein response in cultured myoblasts, underscoring the importance of limiting excess RTL1 during muscle development. Taken together, our findings demonstrate a critical role for AGO2 slicing in skeletal muscle development that is, at least in part, due to its repression of a domesticated retrotransposon.

## INTRODUCTION

Argonaute (AGO) proteins are ancient nucleic acid binding proteins found in all three domains of life. The exquisite sequence specificity of AGO arises from its small RNA or DNA guide, which base-pairs to complementary nucleic acid targets. The ancestral eukaryotic AGO protein is an endonuclease that usually cleaves fully complementary targets at the scissile phosphate bond across from guide nucleotides 10 and 11 (counting from the 5′ end of the guide).^1^ This slicing activity of AGO is thought to have arisen as a host immune response to destroy invading phage, viruses, and transposable elements.^2–6^ However, in mammals, AGOs have largely been repurposed for gene regulation of host transcripts.^7,8^

Mammalian AGOs are primarily loaded with microRNAs (miRNA), 20–24 nucleotide RNAs that guide AGO to bind and repress specific mRNAs. Nearly all miRNA-mediated repression relies on partial complementarity between the seed region (nucleotides 2–8) of a miRNA and its mRNA target, which is not sufficient for endonucleolytic cleavage and instead leads to the recruitment of deadenylating and decapping machinery that accelerates decay and inhibits translation of the mRNA.^7^ This repression is typically modest (i.e. less than 2-fold changes in steady-state mRNA and protein levels for any individual target) but is distributed across dozens to hundreds of targets to broadly shape the transcriptome.

Although partial complementarity is the dominant mode by which AGO–miRNA complexes interact with and regulate targets, the ability to cleave fully complementary targets has been retained by some AGO proteins. In mammals, AGO2 is the main endonuclease,^9–11^ although AGO3 shows some cleavage activity with specific miRNAs and truncated miRNA isoforms.^12,13^ The repression achieved though AGO2-mediated cleavage and subsequent degradation of cleavage fragments by cellular exonucleases is significantly more potent than repression mediated by partial complementarity between a miRNA and target. Indeed, this catalytic activity of AGO2 has been harnessed by researchers and clinicians alike to knock down endogenous transcripts using exogenous short interfering RNAs (siRNAs).^14–18^

Despite the widespread adoption of siRNA technologies, our understanding of the biological functions and substrates regulated by AGO2 slicing is still limited. Mice that express a catalytic-dead AGO2 protein (*Ago2^CD/CD^*) die shortly after birth,^19–21^ indicating a critical role for AGO2 slicing during development. These mice have anemia and impaired red blood cell differentiation caused by defective biogenesis of miR-451 and miR-486-5p, two unusual miRNAs that require AGO2-mediated cleavage to become functional.^19,20,22,23^ Importantly, loss of AGO2 slicing specifically in the hematopoietic compartment or combined loss of miR-451 and miR-486-5p recapitulates the anemia but not the lethality seen in *Ago2^CD/CD^* animals,^20^ indicating that other substrates and cell types must be affected by AGO2 slicing. *Ago2^CD/CD^* mice also have abnormal placental vasculature attributed to increased expression of retrotransposon-like-1 (*Rtl1*), an imprinted protein-coding gene that is regulated in *trans* by several, perfectly complementary miRNAs.^21,24–26^ The extent to which *Rtl1* derepression and vascular defects in the placenta contribute to *Ago2^CD/CD^* embryo lethality is not known. Here, using a combination of histological, molecular, and genetic approaches, we systematically define cell types and substrates that require AGO2 slicing during mouse development.

## RESULTS

### *Ago2^CD/−^* mice recapitulate the lethality and hematopoietic defects of *Ago2^CD/CD^* mice

To expand upon previous studies of AGO2 catalytic activity during mouse development^19–21^ and minimize the potential influence of tightly-linked mutations, we pursued an alternative breeding strategy that relied on two distinct *Ago2* alleles, the previously described catalytic-dead allele (*Ago2^CD^*)^19^ and a knockout allele (*Ago2^−^*), which we generated via cre-mediated recombination of an *Ago2* floxed allele.^27^ *Ago2^CD/−^*mice were recovered at expected Mendelian ratios in utero, but not after birth (Figure 1A, Table S1), similar to reported survival rates for *Ago2^CD/CD^* mice.^19–21^

**Figure 1.**
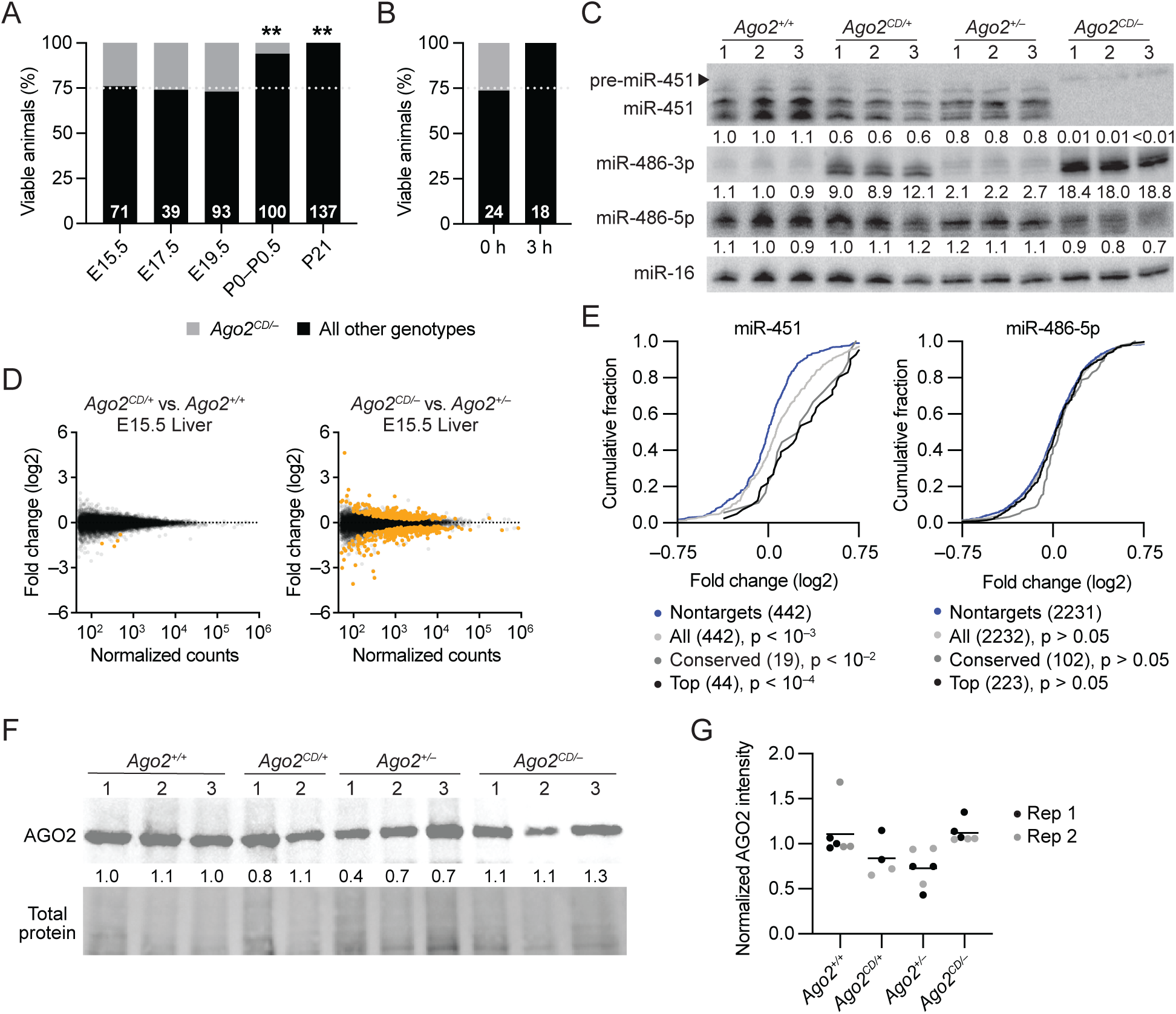
*Ago2^CD/−^*animals recapitulate phenotypes reported in *Ago2^CD/CD^* animals. (A–B) The influence of AGO2 slicing on viability in utero and postnatally. Plotted are the percentage of viable animals at each time point that are either *Ago2^CD/−^* or all other genotypes. The total number of animals evaluated at each time point is indicated. *P* values were calculated for each time point (**, *P* < 0.01; chi-squared test). (C) The influence of AGO2 slicing on miR-451 and miR-486 expression in E15.5 fetal liver. A northern blot measuring miR-451, miR-486-3p, and miR-486-5p levels in fetal liver (3 biological replicates per genotype) is shown. Below each panel, levels of each miRNA are reported relative to levels in the *Ago2^+/+^* fetal liver after normalizing to the level of miR-16. (D) The influence of AGO2 slicing on RNA levels in the E15.5 fetal liver, as determined by RNA-seq. Shown are fold-changes in mean RNA levels for *Ago2^CD/+^*relative to *Ago2^+/+^*, plotted as a function of expression in *Ago2^+/+^* fetal liver (left), and for *Ago2^CD/−^*relative to *Ago2^+/−^*, plotted as a function of expression in *Ago2^+/−^* fetal liver (right). Each dot represents a unique mRNA or noncoding RNA, showing results for all RNAs with at least 50 normalized counts. Orange dots indicate differentially expressed genes as determined by DESeq2 (adjusted *P* value < 0.05). (E) The influence of AGO2 slicing on expression of predicted targets of miR-451 (left) and miR-486-5p (right) in E15.5 fetal liver. Plotted are the cumulative distribution functions of mRNA fold-changes in *Ago2^CD/−^* fetal liver relative to *Ago2^+/−^*fetal liver for all predicted targets (light gray), conserved predicted targets (gray), and top 10% predicted targets (black). The number of genes in each target set is indicated. *P* values were calculated based on a Mann-Whitney test between each target set and a set of 3′ UTR length-matched transcripts with no predicted site for each miRNA family randomly sampled at a 1:1 ratio. Selection of nontarget cohorts for each target set was repeated 21 times and median *P* values are indicated. For simplicity, only nontargets matched to the all targets set are plotted (blue). (F–G) The influence of *Ago2* heterozygosity on AGO2 protein expression in E15.5 fetal liver. A representative western blot measuring AGO2 levels in fetal liver (2–3 biological replicates per genotype) is shown in (F). In each lane, the intensity of AGO2 was normalized to that of Revert total protein stain and reported relative to the average of the *Ago2^+/+^* and *Ago2^CD/+^* samples. The results for two technical replicates (separate gels) are plotted in (G). No significant changes were detected when comparing *Ago2^+/+^* and *Ago2^CD/+^* samples to *Ago2^+/−^*and *Ago2^CD/−^*samples (*P* <0.05; Welch’s t-test). See also Figure S1.

To better define when and why *Ago2^CD/−^* mice died, we collected P0 neonates immediately after birth and assessed their viability over 3 hours in the absence of maternal care. During the observation period, *Ago2^CD/−^* mice were lethargic, showed limited response to gentle physical stimuli, and exhibited agonal breathing indicative of respiratory distress. All *Ago2^CD/−^* mice expired within 3 hours of birth (Figure 1B). *Ago2^CD/−^* mice had similar body weights as littermates (Figure S1A), but were noticeably paler, consistent with the non-lethal anemic phenotypes previously described.^19,20^

The non-lethal anemia in *Ago2^CD/CD^* mice is caused by defective biogenesis of miR-451 and miR-486 in hematopoietic progenitors.^19,20^ To determine whether miRNA expression was similarly affected in *Ago2^CD/−^* mice, we performed small-RNA northern blotting of RNA extracted from fetal liver, the primary site of hematopoiesis during mid-to-late gestation. Indeed, mature miR-451 levels decreased ∼100-fold in E15.5 *Ago2^CD/−^* liver compared to *Ago2^+/−^* liver (Figure 1C), in line with previous reports.^19,20^ Likewise, miR-486-3p, which is normally sliced by AGO2 to facilitate passenger strand removal,^20^ increased ∼18-fold in *Ago2^CD/−^* liver compared to control *Ago2^+/−^* liver (Figure 1C). Intermediate changes to both miR-451 and miR-486-3p were also observed in E15.5 *Ago2^CD/+^* liver, supporting the model that unprocessed miR-451 and miR-486 intermediates accumulate in catalytic-dead AGO2 proteins.^20^

To determine the global impact of miR-451 and miR-486 dysregulation on gene expression, we performed RNA sequencing of E15.5 fetal liver from embryos of all four genotypes generated in our cross. Principal component analysis (PCA) of the 500 most variable genes indicated that the gene expression profiles of *Ago2^CD/−^*fetal livers were more similar to each other and less similar to the profiles from the other three genotypes (Figure S1B). Nonetheless, to minimize potential confounding effects caused by *Ago2* haploinsufficiency, we compared separately mice that expressed either one allele (*Ago2^CD/−^*versus *Ago2^+/−^*) or two alleles (*Ago2^CD/+^* versus *Ago2^+/+^*).

Overall, 676 genes were differentially expressed (adjusted *P* value < 0.05; 347 increased, 329 decreased) in E15.5 *Ago2^CD/−^* liver compared to *Ago2^+/−^* liver, whereas only five genes were differentially expressed in E15.5 *Ago2^CD/+^* liver compared to *Ago2^+/+^*liver (Figure 1D, Table S2–S3). Differentially expressed genes were depleted in gene sets associated with erythrocytes and other hematopoietic cells (Figure S1C), consistent with the known role of miR-451 in red blood cell maturation.^19,20,28–30^

We also compared the fold-changes of predicted miR-451 and miR-486-5p targets to the fold-changes of mRNAs that lacked miR-451 or miR-486-5p sites, with the expectation that higher confidence predictions should correlate with larger effects.^31,32^ Indeed, we observed increased levels of miR-451 targets in E15.5 *Ago2^CD/−^*fetal liver when compared to *Ago2^+/−^* fetal liver (Figure 1E). In contrast, miR-486-5p targets appeared to be minimally affected in *Ago2^CD/−^*fetal liver, in line with prior small-RNA sequencing showing that miR-451 is 3–16-fold more abundant than miR-486-5p in fetal liver erythroblasts.^20,30^ miR-451 and miR-486-5p targets were not significantly increased in *Ago2^CD/+^* liver (Figure S1D), indicating that haploinsufficient processing of miR-451 and miR-486 did not have a substantial impact on the transcriptome. Gene expression and miR-451 targets were also affected, albeit to a lesser extent, in E19.5 *Ago2^CD/−^*fetal liver (Figure S1E–F), likely reflecting the migration of hematopoiesis from liver to bone marrow around the time of birth. In summary, *Ago2^CD/−^*mice phenocopied the lethality and hematopoietic defects observed in *Ago2^CD/CD^*mice and provided an orthogonal strategy to investigate the cell types and substrates that are affected by loss of AGO2 slicing.

### *Ago2* heterozygosity does not reduce AGO2 protein levels

Previous studies have demonstrated that loss of one or more *Ago* genes can be compensated by increased expression of the other paralogs.^33,34^ Thus we wondered whether *Ago2* heterozygosity might also lead to compensatory changes in AGO2 protein levels. To assess AGO2 expression, we first optimized semi-quantitative western blotting across an eight-fold range of total protein input (2.5–20 ug) from mouse fetal liver (Figure S1G). We then quantified AGO2 levels in total protein collected from E15.5 fetal liver of mice with either one or two protein-coding alleles of *Ago2*. No difference in AGO2 expression was detected in *Ago2^CD/−^* and *Ago2^+/−^* fetal liver compared to *Ago2^+/+^* and *Ago2^CD/+^* fetal liver (Figure 1F), consistent with a compensatory response to *Ago2* heterozygosity.

In previous instances of compensation, increased expression of AGO paralogs occurs through protein stabilization or enhanced translation rather than changes in *Ago* mRNA abundance.^33,34^ To determine if *Ago2* haploinsufficiency affected *Ago2* mRNA levels, we reanalyzed our E15.5 fetal liver RNA-seq to compare *Ago2* transcript abundance across all four genotypes. Total *Ago2* levels were decreased 24% in mice with one protein-coding allele compared to mice with two protein-coding alleles, less than the 50% reduction expected for heterozygous animals (Figure S1H). However, our initial analysis focused on all *Ago2* transcripts and we speculated that transcripts arising from the *Ago2^−^* allele may escape nonsense-mediated degradation and contribute to our quantification. To investigate this possibility, we identified sequencing reads that uniquely mapped to the *Ago2^−^* allele, the *Ago2^CD^* allele, or the wild-type allele in each sample. On average, ∼20% of allele-specific reads in *Ago2^+/−^* and *Ago2^CD/−^* fetal liver were derived from the *Ago2^−^* allele (Figure S1I), which when combined with the 24% reduction in total *Ago2* expression, corresponded to a 39% reduction in protein-coding *Ago2* transcripts, indicating that the compensatory response occurred primarily at the protein level.

### AGO2 slicing affects gene expression and substrates in multiple tissues

Having established that *Ago2^CD/−^* mice recapitulate the lethality observed in *Ago2^CD/CD^* mice, we next sought to determine the physiologic basis for this lethality. Common causes of neonatal lethality in the first hours after birth include pulmonary defects, structural defects in the cardiovascular system, musculoskeletal defects affecting respiratory muscles like the diaphragm, neurologic defects affecting muscle innervation and breathing control, and placental defects that affect maturation of the heart and other organs in the embryo proper.^35,36^ To identify a potential role for AGO2 slicing in these tissues, we performed RNA-sequencing on E19.5 heart, lung, placenta, and diaphragm and compared *Ago2^CD/−^* tissues to *Ago2^+/−^*tissues (Figure 2A, Table S2). Of these, *Ago2^CD/−^* diaphragm had the most differentially expressed genes with 420 in total (267 increased, 153 decreased; adjusted *P* value < 0.05). In contrast, only eight genes were differentially expressed when we compared *Ago2^CD/+^* diaphragm to *Ago2^+/+^* diaphragm (Figure S2A). Among the ten most significantly differentially expressed genes in *Ago2^CD/−^* diaphragm (adjusted *P* value < 1 x 10^14^) were three heat shock protein genes (*Hspa1a*, *Hspa1b*, and *Hsph1*), two metabolism-related genes (*Mzpl3* and *Pdk4*), a muscle-specific F-box protein gene (*Fbxo32*) involved in protein degradation and muscle atrophy, and an exapted retrotransposon-like gene (*Rtl1*), all of which were increased at least 3-fold. One of the earliest identified AGO2 cleavage substrates,^24,25^ *Rtl1* was increased in both *Ago2^CD/−^* diaphragm and placenta (Figure 2A, Figure S2B).

**Figure 2:**
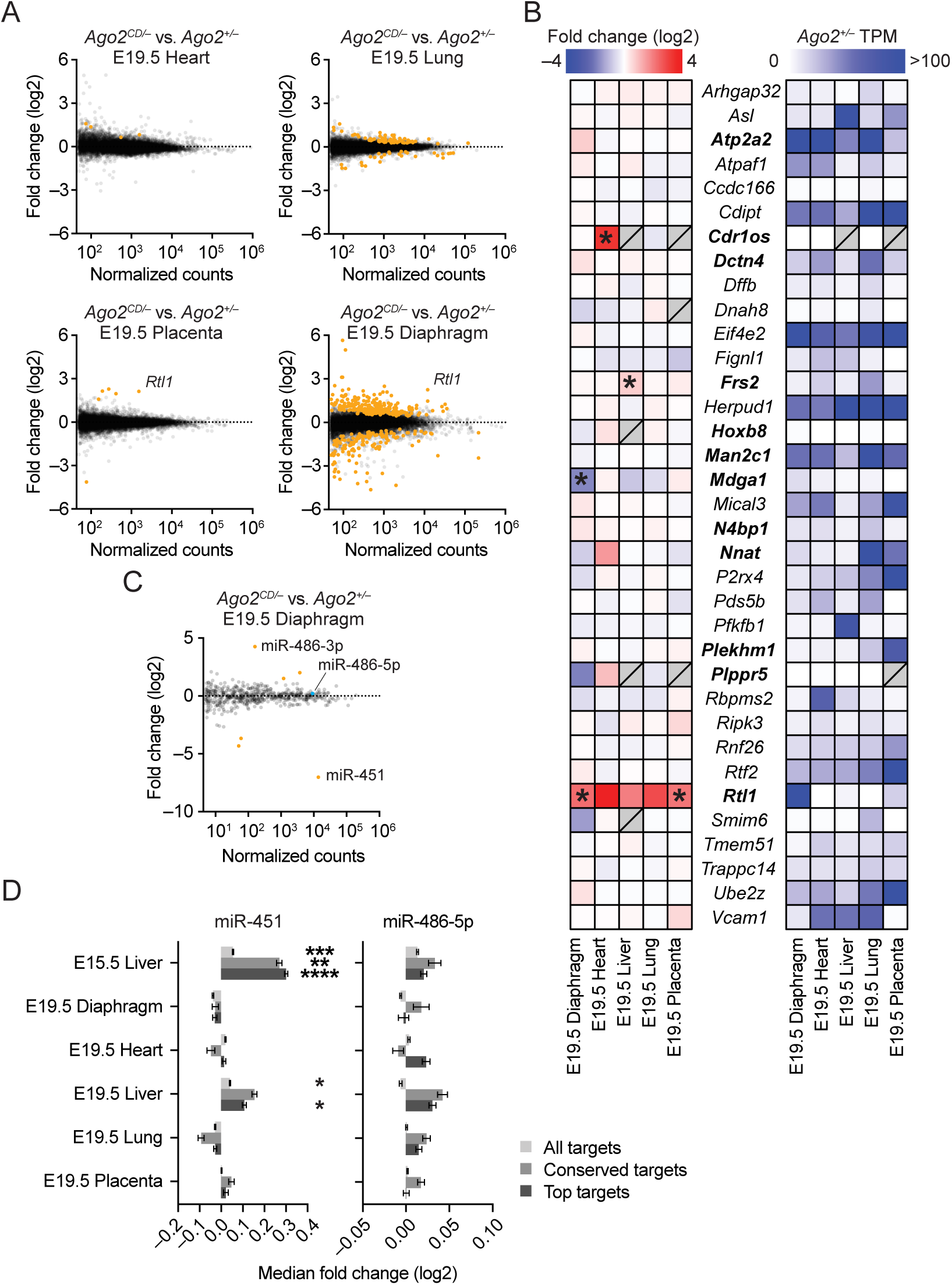
AGO2 endonuclease activity affects gene expression and substrates in multiple tissues. (A) The influence of AGO2 slicing on RNA levels in E19.5 heart, lung, placenta, and diaphragm, as determined by RNA-seq (n = 3–4 per genotype and tissue). Shown are fold-changes in mean RNA levels for *Ago2^CD/−^*tissue relative to *Ago2^+/−^* tissue, plotted as a function of expression in each *Ago2^+/−^* tissue. Each dot represents a unique mRNA or noncoding RNA, showing results for all RNAs with at least 50 normalized counts. Orange dots indicate differentially expressed genes as determined by DESeq2 (adjusted *P* value < 0.05). (B) The influence of AGO2 slicing on known and predicted AGO2 slicing substrates. Heatmaps indicate fold-changes in *Ago2^CD/−^* tissue relative to *Ago2^+/−^* tissue (left) and mean transcripts per million (TPM) in *Ago2^+/−^* tissue (right) for the indicated genes. Bolded gene names denote substrates with miRNA binding sites conserved between mice and humans. Asterisks indicate differentially expressed genes, as determined by DESeq2 (adjusted *P* value < 0.05). Gray squares with slashes indicate contexts in which either fold-changes (left) or TPMs (right) could not be calculated. (C) Influence of AGO2 slicing on miRNA levels in E19.5 diaphragm, as determined by small-RNA-seq (n = 2–3 per genotype). Shown are fold-changes in mean miRNA levels for *Ago2^CD/−^* diaphragm relative to *Ago2^+/−^* diaphragm, plotted as a function of expression in *Ago2^+/−^*diaphragm. Each dot represents a unique miRNA, showing results for all miRNAs with at least five normalized counts per million (CPM). Orange dots indicate differentially expressed miRNAs as determined by DESeq2 (adjusted *P* value < 0.05). (D) Influence of AGO2 slicing on expression of predicted miR-451 and miR-486-5p targets. Plotted are the differences between the median fold change of each set of predicted targets (all, conserved, top) and the median fold change of its matched set of nontargets. This metric of repression was calculated for 21 sampled sets of nontargets, and the mean is displayed (error bars, standard deviation). For each sampling, a *P* value was calculated as described in Figure 1E and the median *P* value is displayed (*, *P* < 0.05; **, *P* < 0.01, ***, *P* < 0.001, ****, *P* < 0.0001). See also Figure S2.

To determine how loss of AGO2 slicing affects other previously reported AGO2 cleavage substrates, we curated a list of bonafide and putative substrates from the literature. In addition to *Rtl1* and the two miRNAs discussed above, 34 AGO2 cleavage substrates have been identified in mouse cells and tissue, 11 of which were also identified in human cells.^37–42^ These substrates were primarily found by degradome sequencing, a method that captures 5′ monophosphate RNA fragments generated by an endonuclease or a 5′–to–3′ exonuclease, combined with computational predictions based on miRNA sequence complementarity.

However, for many substrates, direct evidence that these substrates are regulated by endogenous AGO2 catalytic activity in cells or animals is still lacking. For each substrate, we plotted transcript abundance in *Ago2^+/−^* tissues and fold-changes in *Ago2^CD/−^*tissues compared to *Ago2^+/−^* tissues (Figure 2B, Table S4). Of the 35 putative substrates detected by RNA-seq in at least one tissue, only three were significantly increased: *Frs2* was increased 1.7-fold in E19.5 fetal liver, *Cdr1os* was increased 10-fold in E19.5 heart, and *Rtl1* was increased 4.4-fold in E19.5 placenta and 4.7-fold in E19.5 diaphragm. Increased levels of *Rtl1* were also detected in E19.5 liver, heart, and lung, however, these changes were not significant, likely due to its lower basal expression in those tissues (Figure 2B).

As discussed above, AGO2 slicing is also required for the maturation of miR-451 and miR-486-5p, two miRNAs that are enriched in, but not exclusive to, the hematopoietic compartment.^43^ In fact, miR-486 was first studied in the context of normal and dystrophic muscle^44,45^ and adult *Mir486* knockout mice have smaller muscle fibers and increased frequency of central myonuclei, hallmarks of increased muscle regeneration.^46^ Thus, we wondered if the striking differential gene expression observed in *Ago2^CD/−^* diaphragm (Figure 2A) might be caused by defective processing of miR-486 and/or miR-451. To determine whether loss of AGO2 slicing affects miRNA levels in the diaphragm, we sequenced small RNAs from E19.5 *Ago2^CD/−^* and *Ago2^+/−^*diaphragm. Out of >500 miRNAs detected, only six were differentially expressed (adjusted *P* value < 0.05)(Figure 2C, Table S5). Mirroring what we and others observed with loss of AGO2 slicing in fetal liver, miR-451 levels decreased 128-fold and miR-486-3p levels increased 19-fold in *Ago2^CD/−^* diaphragm. As expected, miR-486-5p levels were unchanged, however, the accumulation of miR-486-3p is expected to prevent miR-486-5p from binding and repressing mRNA targets. In addition, miR-154-3p and miR-485-3p levels increased 4.1- and 2.9-fold, respectively, while miR-137-3p and miR-196a-5p decreased 13- and 20-fold, respectively.

Unlike miR-451 and miR-486, the other miRNAs lack extensive complementarity to their passenger strands and thus are not expected to be suitable AGO2 slicing substrates.

The differential expression of several miRNAs in *Ago2^CD/−^*diaphragm raised the question of whether mRNA targets of these miRNAs might also be affected. To answer this question, we compared fold-changes of predicted targets to fold-changes of mRNAs lacking sites to each miRNA and plotted the difference in median fold-changes. We limited our analyses to miR-451, miR-486-5p, miR-154-3p, and miR-485-3p as the other two differentially expressed miRNAs were lowly expressed in *Ago2^+/−^* diaphragm. Predicted targets of miR-451 and miR-486-5p were not significantly increased in *Ago2^CD/−^*diaphragm (Figure 2D) and predicted targets of miR-154-3p and miR-485-3p were not significantly decreased (Figure S2C), suggesting that these four miRNAs are not abundant enough in muscle to exert measurable repression.

In addition to the substrates discussed above, AGO2 slicing is also involved in the suppression of transposable elements.^28,47–53^ Many TE transcripts are polyadenylated and therefore potentially captured in our RNA-seq data. Using comprehensive TE annotations,^54^ we quantified the expression of TEs in each tissue and compared TE expression in *Ago2^CD/−^* and *Ago2^+/−^*tissues after grouping similar insertions into sub-families. No significant differences in TE expression were detected (Table S6).

Taken together, our RNA-seq and small-RNA-seq analyses identified only a handful of differentially expressed slicing substrates and established that impaired processing of miR-451 and miR-486 did not result in measurable derepression of targets in tissues other than fetal liver. We decided to focus on *Rtl1* because of its robust derepression in both *Ago2^CD/−^* placenta and diaphragm.

### Loss of AGO2 slicing does not affect imprinting at the *Dlk1–Dio3* locus

*Rtl1* is one of three paternally imprinted protein-coding genes in the placental mammal-specific *Dlk1–Dio3* locus. Previous studies have shown that *Rtl1* is regulated in trans by several perfectly complementary miRNAs derived from the maternally imprinted *Rtl1as* transcript.^21,25,26,55^ Because gene expression at imprinted loci is often coordinated, we asked whether increased *Rtl1* expression in *Ago2^CD/−^* tissues was associated with changes to other genes in the *Dlk1–Dio3* locus. In both *Ago2^CD/−^* placenta and *Ago2^CD/−^* diaphragm, no differences were detected in the paternally-expressed protein-coding genes *Dlk1* and *Dio3* or the maternally-expressed noncoding genes *Meg3*, *Rtl1as*, *Rian*, and *Mirg* (Figure S2D), indicating that loss of AGO2 slicing and increased *Rtl1* expression had a negligible effect on imprinting of this locus. In addition, we found no evidence for derepression of other mRNAs targeted by *Rtl1as*-derived miRNAs miR-127, miR-136, miR-431, miR-433 and miR-434. (Figure S2C, E). Overall, our findings are consistent with a previous study showing that increased expression of *Rtl1* due to isolated deletion of *Rtl1as* does not affect the expression of other genes at this locus.^26^

In mice, loss of *Rtl1as* causes early postnatal lethality associated with defects in placental vasculature and skeletal muscle attributed to increased expression of RTL1.^26,55,56^ Of note, this lethality is dependent on genetic background; *Rtl1as*-deficient mice are 100% viable on a mixed 129Sv;C57BL/6 background and < 50% viable after 4–6 backcrosses to C57BL/6.^26^ In humans, loss of *Rtl1as* with or without loss of the other maternally imprinted noncoding RNAs causes Kagami-Ogata syndrome (KOS), which is characterized by bell-shaped thorax, abdominal wall defects including omphalocele, respiratory distress, impaired swallowing and other feeding difficulties, low muscle tone, and placentomegaly.^57–59^ It has been suggested that the abdominal wall, respiratory, and feeding defects in KOS might all arise due to skeletal muscle dysfunction.^56^ These findings highlight the importance of tightly regulating RTL1 expression during mammalian development and raised the possibility that *Ago2^CD/−^*mice might have similar placental and skeletal muscle defects as *Rtl1as*-deficient mice and humans.

### Increased *Rtl1* in *Ago2^CD/−^* placenta does not cause placental defects

During placental development, *Rtl1* is enriched in endothelial cells that line the fetal blood vessels.^26,60^ To determine whether AGO2 slicing represses RTL1 expression in endothelial cells or in other cell types within the placenta, we performed co-immunofluorescence staining using antibodies against mouse RTL1 and endomucin, an endothelial-specific type I membrane protein. This approach did not yield quantitative information about RTL1 levels, but did allow for comparison of RTL1 expression patterns. In both wild-type and *Ago2^CD/−^*placenta, RTL1 expression was mostly detected in endomucin+ endothelial cells (Figure S3A), which was similar to RTL1 expression reported for *Rtl1as*-deficient placenta^26^ and consistent with the model that AGO2 cleavage of *Rtl1* in placenta occurs primarily in fetal endothelium.

In both *Rtl1as*-deficient placenta and *Ago2^CD/CD^*placenta, increased *Rtl1* is associated with placentomegaly and dilated fetal vasculature within the labyrinth zone.^21,26^ Thus, we hypothesized that *Ago2^CD/−^* placental development would be similarly affected. To assess placentomegaly, we measured placental and embryo weights at E19.5. Surprisingly, no differences in placental weight were detected in *Ago2^CD/−^* animals and the fetal-to-placental weight ratio was similar across all four genotypes (Figure 3A, B). To characterize the fetal vasculature of the placenta, we stained sagittal sections of placenta with hematoxylin and eosin or with an antibody against PECAM-1, a marker of endothelial cells, and quantified the relative size of the labyrinth zone, the density of PECAM-1+ vasculature within the labyrinth zone, and the number and diameter of PECAM-1+ vessels (Figure 3B–F). Again, no differences were seen in any of these measurements across the four genotypes, indicating that the fetal vasculature in *Ago2^CD/−^* placenta was not affected, even though *Rtl1* expression was increased in *Ago2^CD/−^*placenta (Figure 2B) by a similar magnitude as in *Ago2^CD/CD^* and *Rtl1as*-deficient placenta.^21,26^ The influence of AGO2 slicing on *Rtl1* expression was not restricted to late gestation; *Rtl1* expression was also increased 4.3-fold in *Ago2^CD/−^* placenta at mid-gestation, when the fetal vascular labyrinth expands to meet the demands of the growing embryo (Figure S3B, C). Taken together, derepression of *Rtl1* in fetal endothelium was not sufficient to cause vascular defects and placentomegaly in *Ago2^CD/−^* placenta.

**Figure 3:**
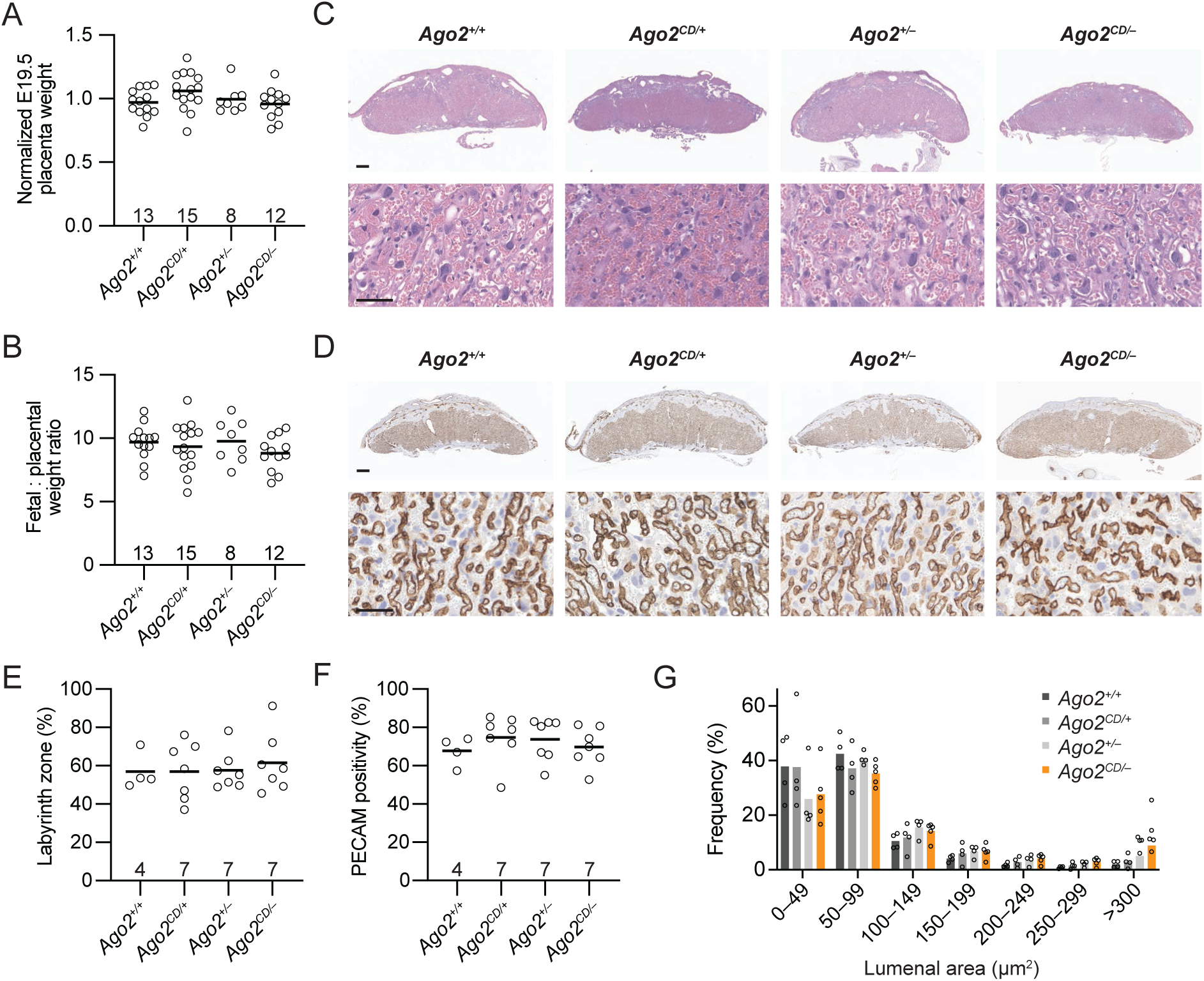
Increased *Rtl1* in *Ago2^CD/−^* placenta does not cause placentomegaly or vascular defects. (A–B) Influence of AGO2 slicing on placental weights. Plotted in (A) are the normalized weights of each placenta at E19.5 (n = 8–15 per genotype). Raw weights were divided by the average weight for each litter to account for litter-to-litter differences. Plotted in (B) are fetal-to-placental weight ratios for each E19.5 animal. No significant changes were detected (*P* <0.05; one-way ANOVA). (C) The influence of AGO2 slicing on E19.5 placental morphology, as determined by hematoxylin and eosin staining. Shown are representative images for each genotype; scale bar, 400 microns (above) and 50 microns (below). (D–G) The influence of AGO2 slicing on E19.5 placental vasculature, as determined by immunohistochemistry for PECAM-1 (brown). Shown in (D) are representative images for each genotype; scale bar, 400 microns (above) and 50 microns (below). The relative area of the labyrinth zone compared to total placental area (E) and the relative area of fetal vasculature, defined by PECAM-1 positivity, within the labyrinth zone (F), are plotted for each genotype (n = 4–7 per genotype). No significant changes were detected (*P* <0.05; one-way ANOVA). Plotted in (G) are the distributions of blood vessel lumenal areas within the fetal vasculature for each genotype (n = 3–5 biological replicates, 365–1281 blood vessels per animal). No significant changes were detected (*P* <0.05; two-tailed Student’s t-test). See also Figure S3.

### *Ago2^CD/−^* skeletal muscle fibers are larger and contain more central nuclei

Finding no differences in *Rtl1*-overexpressing *Ago2^CD/−^* placenta, we turned our attention to the embryo proper. We stained transverse sections of E19.5 torso with hematoxylin and eosin or anti-PECAM-1 antibody and assessed the morphology and vascularization of the heart, lung, and intercostal skeletal muscles. No morphologic differences were detected in *Ago2^CD/−^* heart and lung (Figure S4A–C), consistent with the small number of differentially expressed genes in these tissues (Figure 2A). The fetal vasculature of *Ago2^CD/−^* animals was also similar to littermate controls with no evidence of dilated or leaky vessels (Figure S4B–D).

In contrast, the intercostal muscles of *Ago2^CD/−^* animals were markedly different from that of littermate controls (Figure 4A). When stained with hematoxylin and eosin, *Ago2^CD/−^* intercostal muscle fibers appeared rough and internally disorganized. Quantitatively, *Ago2^CD/−^*intercostal muscle fibers had a broader distribution of sizes and a median cross-sectional area of 170.5 µm^2^, which was 1.9–2.5-fold greater than the median cross-sectional area of muscle fibers from other genotypes (Figure 4B, C). In addition, 16% of *Ago2^CD/−^* muscle fibers had central nuclei, a 1.6–7.3-fold increase compared to fibers from other genotypes (Figure 4D). Central myonuclei are normal during embryonic development, but by E19.5, most myonuclei have migrated to the periphery of muscle cells.^61^ We wondered if the increase in central nuclei might reflect a delay in *Ago2^CD/−^* muscle maturation and investigated this possibility by comparing the expression of embryonic and fetal/neonatal myosin heavy chain genes in our samples. No significant differences were detected in *Ago2^CD/−^*diaphragm (Table S2), arguing against a defect in muscle maturation.

**Figure 4:**
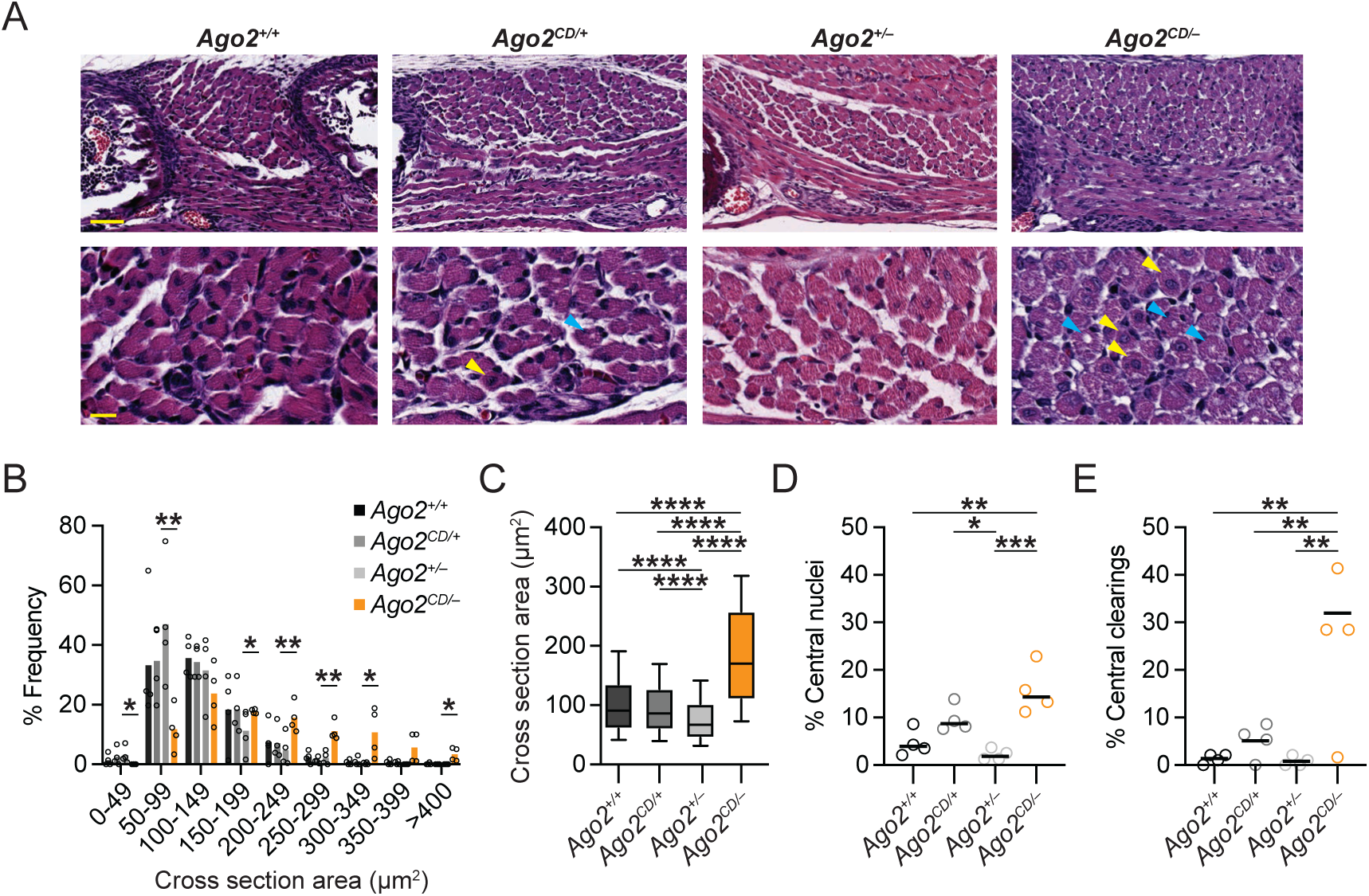
*Ago2^CD/−^* skeletal muscle fibers are larger and contain more central nuclei. (A–E) The influence of AGO2 slicing on skeletal muscle morphology, as determined by hematoxylin and eosin staining. Shown in (A) are representative images of E19.5 intercostal muscle for each genotype; scale bar, 50 microns (above) and 20 microns (below). Yellow arrowheads indicate central nuclei, and the blue arrowheads indicate central clearings within muscle fibers. Plotted in (B) are the distributions of muscle fiber cross sectional area (CSA) in E19.5 intercostal muscles (n = 4 per genotype; 198–679 fibers per animal; *, *P* < 0.05, **, *P* < 0.01; two-tailed Student’s t-test). Plotted in (C) are the median, 25^th^/75^th^ percentile (box), and 10^th^/90^th^ percentile (whiskers) CSAs of all fibers for each genotype (***, *P* < 0.001; ****, *P* < 0.0001; one-way ANOVA with Kruskal-Wallis test). The percentage of fibers with central nuclei (D) and percentage of fibers with central clearings (E) are plotted for each genotype (n = 4 per genotype; *, *P* < 0.05; **, *P* < 0.01; ***, *P* < 0.001; one-way ANOVA with Tukey’s multiple comparisons test). See also Figure S4.

*Ago2^CD/−^* muscle fibers also had an increased frequency of central clearings, areas within the fiber that had minimal hematoxylin and eosin staining and resembled vacuoles associated with some congenital myopathies (Figure 4E).^62^ The central clearings did not stain with trichrome or Periodic Acid-Schiff (Figure S4E, F), indicating that the clearings were not associated with deposits of collagen or glycogen, respectively. We also found no evidence of increased muscle fibrosis in *Ago2^CD/−^* animals (Figure S4E). Taken together, the histologic changes detected in *Ago2^CD/−^* intercostal muscles are consistent with a developmental defect in skeletal muscle maturation and raise the possibility that AGO2 slicing in skeletal muscle is required for postnatal viability.

### AGO2 slicing is required in skeletal muscle for postnatal viability

To test the importance of AGO2 slicing in skeletal muscle, we used a cre recombinase-based strategy to generate mice that express only catalytic-dead AGO2 in skeletal muscle and both wild-type and catalytic-dead AGO2 in all other tissues. This strategy was previously used to demonstrate cell-autonomous roles for AGO2 slicing in hematopoietic cells and female oocytes.^20,52^ Here, we bred homozygous *Ago2^fl/fl^* mice to *Ago2^CD/+^* mice that also carried a *cre* allele expressed from the skeletal muscle-specific human *ACTA1* promoter.^63^ This breeding generated four genotypes and we compared the viability of the *ACTA1-cre+*; *Ago2^CD/fl^* mice to the viability of the other three genotypes (Figure 5A). Of the 84 animals genotyped at weaning, only one *ACTA1-cre+*; *Ago2^CD/fl^* animal was detected; this animal was euthanized because it was too small to wean and genotyping of DNA extracted from the diaphragm showed incomplete excision of the *Ago2* floxed allele. Similar lethality was observed when *cre* was expressed ubiquitously from a *CMV* promoter. For both *ACTA1-cre+*; *Ago2^CD/fl^*and *CMV-cre+*; *Ago2^CD/fl^* mice, lethality occurred shortly after birth (Table S1), recapitulating the lethality seen in *Ago2^CD/−^* mice (Figure 1A). In contrast, *VEC-cre+*; *Ago2^CD/fl^*, *Mlc2v-cre+*; *Ago2^CD/fl^*, and *Actl6b-cre+*; *Ago2^CD/fl^*mice were recovered at expected Mendelian ratios, indicating that AGO2 slicing in endothelium (including fetal endothelium of the placenta), cardiomyocytes, and neurons, respectively, was dispensable for normal development (Figure 5A).

**Figure 5:**
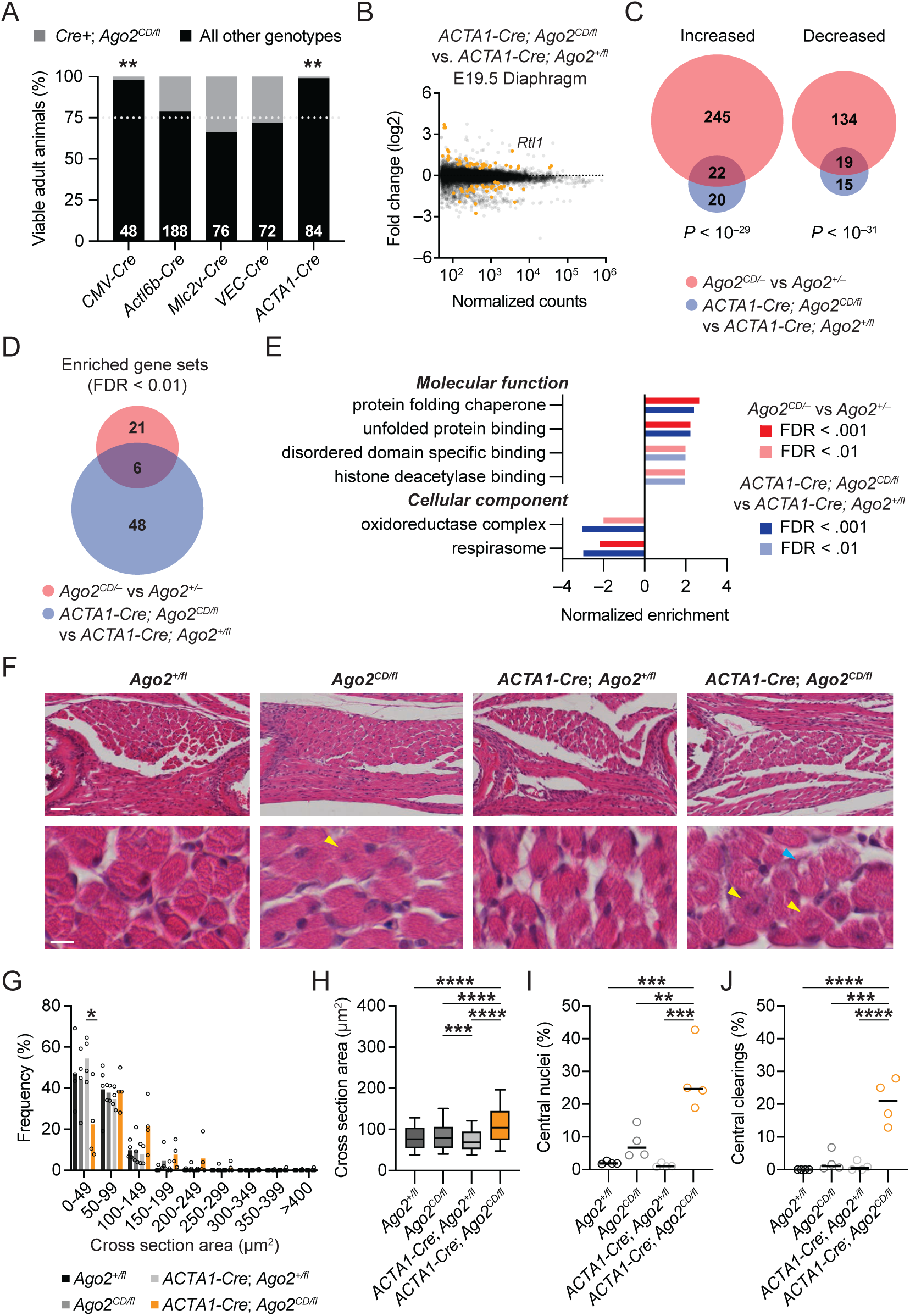
Loss of AGO2 endonuclease activity in skeletal muscle causes muscle defects and perinatal lethality. (A) The influence of cell type-specific loss of AGO2 slicing on postnatal viability. Plotted are the percentage of viable animals at weaning age that were either *Cre+*; *Ago2^CD/fl^* or any other genotype. The total number of animals evaluated and statistical significance for each *Cre*-expressing mouse line is indicated (**, *P* < 0.01; chi-squared test). (B) The influence of skeletal muscle-specific loss of AGO2 slicing on RNA levels in E19.5 diaphragm (n = 2–3 per genotype). Shown are fold-changes in mean RNA levels for *ACTA1-Cre*; *Ago2^CD/fl^* relative to *ACTA1-Cre*; *Ago2^+/fl^*, plotted as a function of expression in *ACTA1-Cre*; *Ago2^+/fl^* diaphragm. Each dot represents a unique mRNA or noncoding RNA, showing results for all RNAs with at least 50 normalized counts. Orange dots indicate differentially expressed genes as determined by DESeq2 (adjusted *P* value < 0.05). (C) The overlap of differentially expressed genes (adjusted *P* value < 0.05) in E19.5 *Ago2^CD/−^* and *ACTA1-Cre*; *Ago2^CD/fl^* diaphragm. The significance of the overlap is indicated (Fisher’s exact test). (D–E) Gene set enrichment analysis for E19.5 *Ago2^CD/−^* and *ACTA1-Cre*; *Ago2^CD/fl^* diaphragm. Shown in (D), the overlap of significantly enriched gene sets (FDR < 0.01) associated with molecular function and cellular component, as determined by WebGestalt. Plotted in (E) are normalized enrichment scores for the six shared gene sets. (F–J) The influence of skeletal muscle-specific loss of AGO2 slicing on skeletal muscle morphology, as determined by hematoxylin and eosin staining. Shown in (F) are representative images of E19.5 intercostal muscle for each genotype; scale bar, 50 microns (above) and 20 microns (below). Yellow arrowheads indicate central nuclei, and the blue arrowheads indicate central clearings within muscle fibers. Plotted in (G) are the distributions of muscle fiber cross sectional area (CSA) in E19.5 intercostal muscles (n = 4 per genotype, 93–303 fibers per animal; *, *P* < 0.05; two-tailed Student’s t-test). Plotted in (H) are the median, 25^th^/75^th^ percentile (box), and 10^th^/90^th^ percentile (whiskers) CSAs of all fibers for each genotype (***, *P* < 0.001; ****, *P* < 0.0001; one-way ANOVA with Kruskal-Wallis test). The percentage of fibers with central nuclei (I) and percentage of fibers with central clearings (J) are plotted for each genotype (n = 4 per genotype; **, *P* < 0.01; ***, *P* < 0.001, ****, *P* < 0.0001; one-way ANOVA with Tukey’s multiple comparisons test). See also Figure S5.

### *Rtl1* is increased in *ACTA1-cre+*; *Ago2^CD/fl^* diaphragm

To determine whether the transcriptomic changes seen in *Ago2^CD/−^*diaphragm were caused by loss of AGO2 slicing specifically in skeletal muscle, we performed RNA-sequencing on E19.5 diaphragm from *ACTA1-cre+*; *Ago2^+/fl^*and *ACTA1-cre+*; *Ago2^CD/fl^* mice (Figure 5B, Table S7). Overall, 78 genes were differentially expressed (adjusted *P* value < 0.05; 44 increased, 34 decreased) in *ACTA1-cre+*; *Ago2^CD/fl^*diaphragm. Similar to what we observed in *Ago2^CD/−^*diaphragm, *Rtl1* was increased 3.5-fold in *ACTA1-cre+*; *Ago2^CD/fl^*diaphragm. We confirmed the increased expression of *Rtl1* by RT-qPCR (Figure S5A) with primer pairs that recognize the skeletal muscle-enriched isoform of *Rtl1* (Figure S2B). Next, we compared the 78 differentially expressed genes in *ACTA1-cre+*; *Ago2^CD/fl^* diaphragm to the 420 differentially expressed genes in *Ago2^CD/−^* diaphragm (Figure 2A); a significant overlap was observed for both upregulated and downregulated genes (Figure 5C). To identify additional pathways affected by loss of AGO2 slicing in muscle, we performed gene set enrichment analysis for molecular functions and cellular components. In total, 27 and 54 gene sets were significantly enriched/depleted in *Ago2^CD/−^*and *ACTA1-cre+*; *Ago2^CD/fl^* diaphragm, respectively, with six of these gene sets significantly enriched/depleted in both comparisons (FDR < 0.01)(Figure 5D, Table S8). The most significantly enriched gene sets in both *Ago2^CD/−^*and *ACTA1-cre+*; *Ago2^CD/fl^* diaphragm were related to chaperones and unfolded protein binding, whereas the most significantly depleted gene sets were related to mitochondrial function and the electron transport chain (Figure 5E). Notably, gene sets associated with the unfolded protein response or the integrated stress response were not significantly enriched in either *Ago2^CD/−^* and *ACTA1-cre+*; *Ago2^CD/fl^* diaphragm.

To determine which transcription factors might be driving the gene expression changes observed in *Ago2^CD/−^* and *ACTA1-cre+*; *Ago2^CD/fl^* diaphragm, we performed overrepresentation analysis using the genes that were either significantly increased or decreased (adjusted *P* value < 0.05) from each comparison and the 2022 ChIP-X Enrichment Analysis database of transcription factor targets from Enrichr.^64,65^ The only transcription factor whose targets were enriched in both sets of upregulated genes was heat shock factor 1 (HSF1), the master regulator of cytosolic proteostasis (odds ratio of 5.2, adjusted *P* value < 1 x 10^−11^ in *Ago2^CD/−^*diaphragm; odds ratio of 7.3, adjusted *P* value < 0.02 in *ACTA1-cre+*; *Ago2^CD/fl^* diaphragm). No transcription factor targets were enriched in either set of downregulated genes.

### *ACTA1-cre+*; *Ago2^CD/fl^* skeletal muscle fibers are larger and contain more central nuclei

To determine whether the muscle defects seen in *Ago2^CD/−^*mice were caused by loss of AGO2 slicing specifically in skeletal muscle, we performed a similar histologic assessment on *ACTA1-cre+*; *Ago2^CD/fl^*mice and littermate controls. Like *Ago2^CD/−^* intercostal muscle fibers, *ACTA1-cre+*; *Ago2^CD/fl^* intercostal muscle fibers appeared rough and internally disorganized (Figure 5F). *ACTA1-cre+*; *Ago2^CD/fl^* intercostal muscle fibers had a median cross-sectional area of 104.0 µm^2^, which was 1.3–1.5-fold greater than the median cross-sectional area of fibers from other genotypes (Figure 5G–H). In addition, *ACTA1-cre+*; *Ago2^CD/fl^* muscle fibers had an increased frequency of central nuclei and central clearings when compared to fibers from other genotypes (Figure 5I–J). Taken together, skeletal muscle-specific loss of AGO2 slicing recapitulated the key histologic and molecular changes observed with global loss of AGO2 slicing, indicating that these changes were largely cell autonomous.

### RTL1 overexpression dramatically remodels the myoblast transcriptome

Given the shared derepression of *Rtl1* in *Ago2^CD/−^*and *ACTA1-cre+*; *Ago2^CD/fl^* diaphragm and previous reports in both humans and mice linking increased expression of RTL1 to muscle weakness and dysfunction,^56,58,66^ we posited that increased RTL1 was a major driver of the transcriptomic changes observed in *Ago2^CD/−^* and *ACTA1-cre+*; *Ago2^CD/fl^*diaphragm. To test this, we generated polyclonal C2C12 mouse myoblast cell lines that express either mouse *Rtl1*, human *RTL1*, or *Halo* under the control of a doxycycline-inducible promoter (Figure 6A). Of note, no differences in cell viability were observed for untreated and doxycycline-treated cell lines. We measured RTL1 levels by Western blot using total protein extracted from C2C12 cells and an antibody raised against mouse RTL1 (Figure 6B). Endogenous RTL1 was not detected in *Halo* control cells. RTL1 was also not detected in untreated *Rtl1*/*RTL1* cells, indicating that our dox-inducible system had minimal leakiness. After 24 hours of doxycycline treatment, robust expression of both mouse and human RTL1 proteins were detected; these proteins each migrated as a single dominant band at the expected size. To determine the subcellular localization of RTL1, we performed confocal microscopy on live *Rtl1-Halo*-expressing cells stained with membrane-permeable Halo ligand (Figure S6A). Mouse RTL1-Halo was detected diffusely throughout the cytosol with no evidence of aggregation.

**Figure 6:**
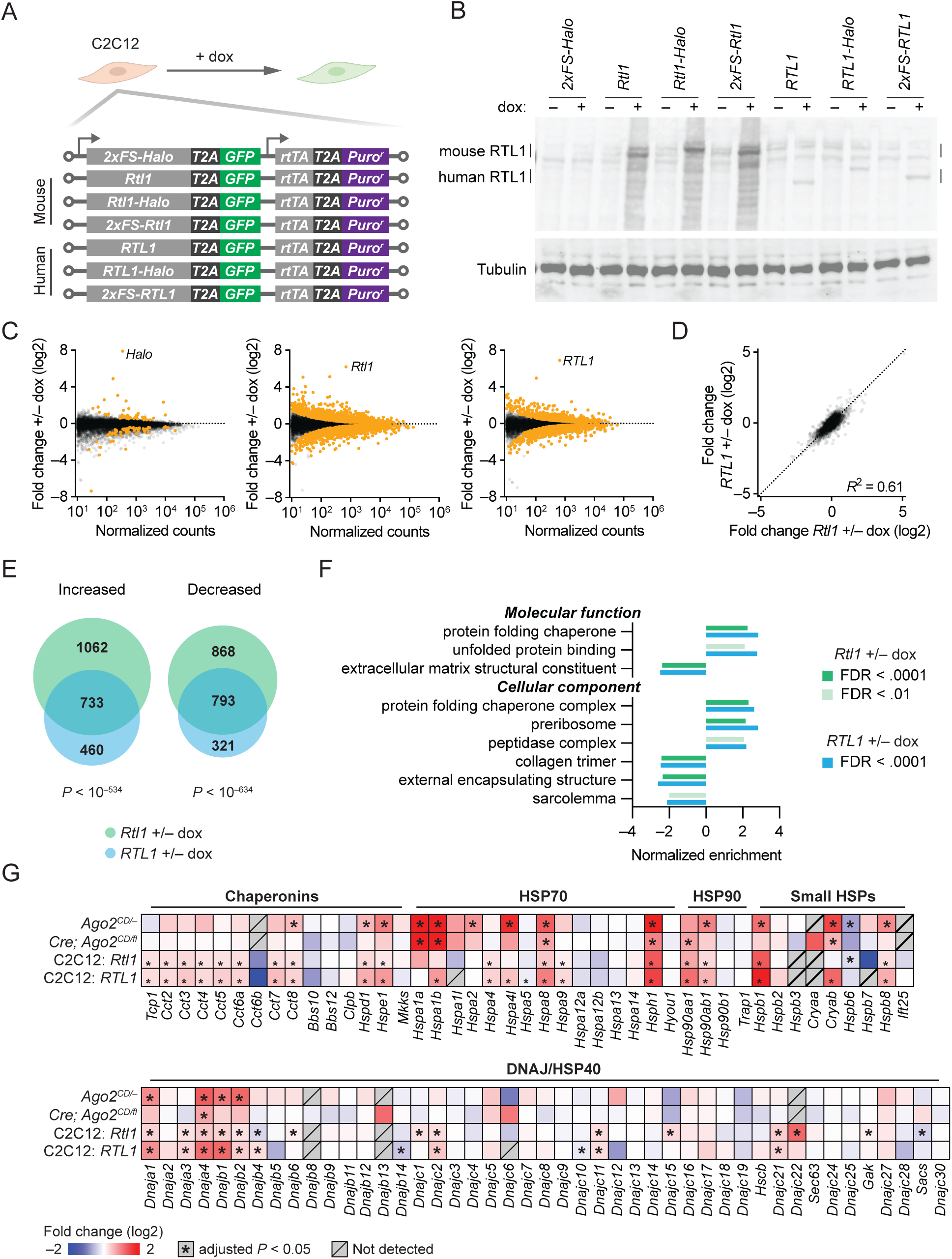
RTL1 expression induces a heat shock response in muscle cells. (A) Engineered C2C12 cell lines used to determine the influence of RTL1 on myoblast gene expression. Polyclonal lines were generated by stably integrating one of seven doxycycline-inducible expression cassettes: a 2xFLAG/Strep-tagged Halo control, three versions of mouse RTL1 (untagged, N-terminal 2xFLAG/Strep-tagged, and C-terminal Halo-tagged), and three versions of human RTL1 (untagged, N-terminal 2xFLAG/Strep-tagged, and and C-terminal Halo-tagged). All expression cassettes also contain T2a-GFP. (B) A representative western blot measuring RTL1 levels in untreated and doxycycline-treated cell lines. The estimated mass of untagged mouse and human RTL1 are ∼199 kDa and ∼155 kDa, respectively; the C-terminal Halo tag adds ∼34 kDa to each. (C) The influence of RTL1 on RNA levels in C2C12 cells, as determined by RNA-seq (n = 3 biological replicates per condition). Shown are fold-changes in mean RNA levels for doxycycline-treated *Halo* (left), mouse *Rtl1* (middle), and human *RTL1* (right) cell lines relative to untreated lines, plotted as a function of expression in untreated lines. Each dot represents a unique mRNA or noncoding RNA, showing results for all RNAs with at least 50 normalized counts. Orange dots indicate differentially expressed genes as determined by DESeq2 (adjusted *P* value < 0.05). (D) Correlation of fold-changes associated with increased RTL1 in C2C12 cells. Plotted are the fold-changes in *Rtl1*-expressing cells versus the fold-changes in *RTL1*-expressing cells for all genes with at least 100 mean normalized counts in the untreated condition. The correlation coefficient (Pearson *R*^2^) is indicated. (E) The overlap of differentially expressed genes (adjusted *P* value < 0.05) in *Rtl1/RTL1*-expressing cells after removing 60 genes that were also differentially expressed in *Halo*-expressing cells. The significance of the overlap is indicated (Fisher’s exact test). (F) Gene set enrichment analysis for doxycycline-treated mouse *Rtl1* and human *RTL1* cell lines, as determined by WebGestalt. Plotted are normalized enrichment scores for the nine significantly enriched gene sets (FDR < 0.01) associated with molecular function and cellular component that were identified in both comparisons. (G) The influence of AGO2 slicing and RTL1 overexpression on heat shock protein genes. Heatmaps indicate fold-changes in *Ago2^CD/−^* or *ACTA1-cre*; *Ago2^CD/fl^*diaphragm compared to their respective controls or in *Rtl1/RTL1*-expressing cells treated with doxycycline compared to untreated cells. Asterisks indicate differentially expressed genes, as determined by DESeq2 (adjusted *P* value < 0.05). Gray squares with slashes indicate contexts in which fold-changes could not be calculated.

To determine the impact of RTL1 expression on the C2C12 transcriptome, we performed RNA-sequencing on doxycycline-treated and untreated cell lines (Figure 6C). RTL1 expression resulted in broad remodeling of RNA levels in C2C12 myoblasts; thousands of genes were differentially expressed in cells expressing mouse *Rtl1* (adjusted *P* value < 0.05; 1824 increased, 1689 decreased) or human *RTL1* (adjusted *P* value < 0.05; 1219 increased, 1135 decreased), whereas only 115 genes were differentially expressed in cells expressing the *Halo* control (adjusted *P* value < 0.05; 59 increased, 56 decreased)(Figure 6C, Table S9). The levels of *Rtl1/RTL1* in doxycycline-treated C2C12 myoblasts were similar to the levels observed with loss of AGO2 slicing in mouse diaphragm: 554.31 ± 168.39 TPM (mean and standard deviation) in *Rtl1*-expressing cells, 1850.12 ± 151 TPM in *RTL1*-expressing cells, 729.45 ± 200.96 TPM in *Ago2^CD/−^* diaphragm, and 508.46 ± 66.93 TPMs in *ACTA1-cre+*; *Ago2^CD/fl^*diaphragm (Table S4). The fold-changes in *Rtl1*-expressing cells also correlated with the fold-changes in *RTL1*-expressing cells (Pearson *R*^2^ = 0.61) and a significant overlap was noted for both upregulated and downregulated genes, consistent with a conserved regulatory activity (Figure 6D, E). In contrast, the fold-changes in *Rtl1*- or *RTL1*-expressing cells showed minimal correlation with the fold-changes in *Halo*-expressing cells (Pearson *R*^2^ = 0.04 and 0.06, respectively). To identify the pathways most affected by RTL1 expression, we performed gene set enrichment analysis for molecular functions and cellular components. In total, 11 and 31 gene sets were significantly enriched/depleted in *Rtl1*- and *RTL1*-expressing cells, respectively, with nine of these gene sets significantly enriched/depleted in both comparisons (FDR < 0.01)(Figure 6F, Table S8). Similar to what we observed in *Ago2^CD/−^* and *ACTA1-cre+*; *Ago2^CD/fl^*diaphragm, the most significantly enriched gene sets in *Rtl1*- and *RTL1*-expressing cells were related to chaperones and unfolded protein binding, whereas the most significantly depleted gene sets were related to extracellular matrix and muscle cell membranes. Again, genes that are normally induced by the unfolded protein response or integrated stress response were unchanged, consistent with an isolated induction of protein folding chaperone genes. Overrepresentation analysis of transcription factor targets demonstrated strong enrichment of HSF1, MYC, and CREB/CREM targets among the genes that significantly increased with either mouse or human RTL1 overexpression and strong enrichment of Polycomb repressive complex 2 targets among the genes that significantly decreased with either mouse or human RTL1 overexpression (Figure S6B, C). We also compared the fold-changes of heat shock protein genes, many of which are direct targets of HSF1, across all four RNA-seq analyses, two from the diaphragm and two from C2C12 myoblasts (Figure 6G). Of the 88 heat shock protein genes detected in at least one analysis, 12 genes representing all five major classes of heat shock protein were significantly increased in three analyses and three genes, *Dnaja4*, *Hspa8*, and *Hsph1*, were significantly increased in all four analyses. Overall, these results support a model in which increased expression of RTL1, both *in vivo* and *in vitro*, induces a broad HSF1-driven heat shock response, which although presumably protective, is not sufficient to prevent muscle defects in vivo.

## DISCUSSION

While the influence of AGO2 slicing has been reported for some fetal and postnatal tissues, the critical cell-types and substrates that explain the postnatal lethality associated with loss of AGO2 slicing have remained elusive. Here, we performed systematic phenotyping and molecular profiling of five fetal tissues and genetic ablation of AGO2 slicing in four distinct cell lineages. Our work definitively demonstrates that AGO2 slicing is required in skeletal muscle for proper muscle development and postnatal survival, and implicates excess *Rtl1* as a major driver of the muscle defects associated with loss of AGO2 slicing.

### Primary defect occurs in skeletal muscle

Although global loss of AGO2 slicing affected gene expression in multiple embryonic tissues, the changes detected in skeletal muscle were the most striking. More than 400 genes were differentially expressed in *Ago2^CD/−^*diaphragm (Figure 2A). These gene expression changes were associated with prominent changes in skeletal muscle morphology *in vivo*. *Ago2^CD/−^*intercostal muscle fibers had a 2-fold increase in median cross-sectional area, a 2–7-fold increase in central nuclei, and a 6–40-fold increase in central clearings compared to fibers from other genotypes (Figure 4A–E). We also noted a qualitative difference in the texture of *Ago2^CD/−^*muscle fibers, which might reflect structural damage or impaired interactions with the extracellular matrix. Importantly, mice with skeletal muscle-specific loss of AGO2 slicing largely recapitulated the molecular and histologic changes, as well as the postnatal lethality, associated with global loss of AGO2 slicing (Figure 5A–J).

The muscle phenotypes in *Ago2^CD/−^* and *ACTA1-cre+*; *Ago2^CD/fl^* animals overlap with but are distinct from the pathologic changes seen in muscle diseases caused by impaired autophagy and/or lysosomal function such as centronuclear myopathy and Pompe’s disease.^62,67,68^ For instance, central nuclei are the eponymous feature of centronuclear myopathies, and sometimes accompanied by perinuclear glycogen deposits. However, the fibers in centronuclear myopathies tend to be smaller in size, whereas *Ago2^CD/−^* and *ACTA1-cre+*; *Ago2^CD/fl^* fibers were significantly larger. Central clearings, without central nuclei, are common in lysosomal storage disorders like Pompe’s disease. These clearing represent an accumulation of lysosomes filled with glycogen, which stains poorly with hematoxylin and eosin but can be detected by Periodic Acid-Schiff staining. We did not detect PAS-positive material in the central clearings of *Ago2^CD/−^*fibers (Figure S4F), indicating that these clearings likely arise through a different mechanism. Whether the defects seen in *Ago2^CD/−^* and *ACTA1-cre+*; *Ago2^CD/fl^*muscle fibers are the cause or consequence of muscle dysfunction remains to be determined.

### Increased *Rtl1* is a driver of the muscle pathology and lethality in *Ago2^CD/−^* mice

Three known slicing substrates were affected by loss of AGO2 slicing in the diaphragm; *Rtl1* increased 4.7-fold, miR-451 decreased 128-fold, and miR-486-3p increased 19-fold. However, we were unable to detect derepression of miR-451 and miR-486-5p targets, indicating that the changes in miRNA levels have minimal influence on the skeletal muscle transcriptome. Instead, multiple lines of evidence point to *Rtl1* as a critical mediator of the muscle pathology and lethality observed in *Ago2^CD/−^* mice: 1) *Rtl1as* deficiency and consequent derepression of *Rtl1* causes widespread pathologic changes in neonatal skeletal muscle, including increased muscle fiber size and central nuclei,^56^ that mirror our findings in *Ago2^CD/−^* muscle, 2) *Rtl1as* deficiency in humans causes Kagami-Ogata syndrome, which is characterized by poor feeding, respiratory distress, and other symptoms ascribed to muscle weakness,^57,58^ and 3) ectopic expression of miRNA-resistant *Rtl1* in embryonic skeletal muscle causes impaired diaphragm development and postnatal lethality, although diaphragm muscle fibers are reportedly smaller in size.^66^ Additional support for an *Rtl1*-centric interpretation of our findings comes from gene expression profiling. We find that RTL1 overexpression in C2C12 myoblasts induces gene expression programs related to chaperones and unfolded protein binding, the same programs that are enriched in *Ago2^CD/−^* and *ACTA1-cre+*; *Ago2^CD/fl^* diaphragm (Figure 5D, 6F). Increased expression of chaperones and other HSF1 targets, together with evidence that RTL1 co-immunoprecipitates with heat-shock proteins,^66^ point to a coordinated cellular response to excess cytosolic RTL1, which because of its large viral-like capsid domain,^69^ may require chaperones to promote folding and/or prevent oligomerization. The features or function(s) of RTL1 that induce a heat shock response appear to be conserved as expression of human RTL or mouse RTL1 causes similar remodeling of the C2C12 transcriptome.

Even though *Rtl1* is a critical mediator of *Ago2^CD/−^*muscle pathology and postnatal lethality, we suspect that increased *Rtl1* is not sufficient to confer the full spectrum of phenotypes caused by loss of AGO2 slicing in skeletal muscle. We make this inference based on differential penetrance of postnatal lethality in *Rtl1as*-deficient and *Ago2^CD/−^*animals. *Rtl1as*-deficient mice are 100% viable on a mixed 50% C57BL/6; 50% 129/Sv background and exhibit partially penetrant lethality after 4–6 backcrosses into C57BL/6.^26^ Even with 10 backcrosses into C57BL/6, some viable *Rtl1as*-deficient mice are recovered.^70^ On the other hand, loss of AGO2 slicing is 100% lethal in our mice, which are primarily on a C57BL/6 background (96.3–98.5% C57BL6/J and C57BL/6NCrl per Transnetyx SNP analysis, which is equivalent to 5–6 backcrosses). To determine if background affected phenotypic penetrance in *Ago2^CD/−^* mice, we crossed *Ago2^CD/+^* and *Ago2^+/−^* mice to wild-type 129X1/SvJ mice and then mated heterozygous F1 animals, generating *Ago2^CD/−^* progeny and littermates that were 50% 129X1/SvJ; 50% C57BL/6. No *Ago2^CD/−^* animals were detected at weaning (Table S1), indicating that the lethality associated with loss of AGO2 slicing is more penetrant than the lethality associated with *Rtl1as* deficiency. Similar results were obtained when we backcrossed to DBA/2J (Table S1). We hypothesize that the differential penetrance may be due to regulation of one or more unidentified slicing substrates in skeletal muscle.

### Increased *Rtl1* in *Ago2^CD/−^* placenta is not sufficient to cause placental vascular defects

One surprise from our study was the lack of placentomegaly and vascular defects in the *Ago2^CD/−^* placenta, even though *Rtl1* levels were increased 4-fold in both E15.5 and E19.5 placenta (Figure 2A, S3B). Similar magnitude fold-changes in *Rtl1* were reported for *Rtl1as*-deficient (*Rtl1^MatKO/+^*) and *Ago2^CD/CD^* placenta, both of which had placentomegaly and vascular defects,^21,26^ indicating that increased *Rtl1* is required, but not sufficient, to cause vascular defects in the placenta. We did detect increased fold-changes in some endothelial-associated genes, especially in E15.5 *Ago2^CD/−^* placenta, however, these fold-changes did not have a measurable impact on the amount or morphology of placental vasculature in E19.5 placenta (Table S2, Figure 3B–F).

### Most AGO2 slicing substrates are not derepressed

Another surprise from our study was the lack of derepression for most predicted or established AGO2 slicing substrates. Of the 35 substrates we examined, only three were significantly increased in any of our RNA-seq libraries; *Frs2* increased in E15.5 and E19.5 fetal liver, *Cdr1os* increased in E19.5 heart, and *Rtl1* increased in E15.5 placenta, E19.5 placenta, and E19.5 diaphragm. At least one slicing substrate was identified in each tissue, with the exception of E19.5 lung, indicating that AGO2 expression is not limiting. The lack of derepression for other substrates may be due to low expression of the substrate (although more than half were expressed at >40 TPM in at least one tissue), low expression of the miRNA that binds the substrate, or perhaps, false positive identification of substrates in previous studies. In theory, the loss of AGO2 slicing could also be partially compensated by canonical microRNA-mediated repression, as the catalytic-dead AGO2 can still bind the substrate and recruit deadenylation and decapping machinery. In that respect, the levels of *Rtl1* might be expected to increase more in *Rtl1as*-deficient mice, which lack the perfectly complementary miRNAs to *Rtl1*, when compared to *Ago2^CD/−^*mice. Although we are unable to make that comparison directly, it doesn’t appear to be the case in placenta or diaphragm.^26,56^ The miRNA concentration needed to repress targets via deadenylation is presumably much higher than the concentration needed for efficient slicing.^71,72^

### Limitations

We focused our investigation on respiratory muscles because of their critical importance for postnatal survival and our initial observations of impaired breathing in P0 neonates. The diaphragm is the primary muscle for respiration and intercostal muscles help form and move the chest wall, which is particularly important during deep inhalation needed to open the airway after birth. However, it is possible that the influence of AGO2 slicing is limited to specific fiber types or muscle groups. Another potential limitation of our work is the comparability of *in vivo* and i*n vitro* gene expression programs. While we attempted to minimize the difference by expressing *Rtl1*/*RTL1* in C2C12 cells at levels similar to what we observed in *Ago2^CD/−^* and *ACTA1-cre+*; *Ago2^CD/fl^*, we acknowledge the many differences between C2C12 myoblasts and mature muscle fibers, as well as differences in metabolism, extracellular matrix, and microenvironment that are not accounted for *in vitro*. Finally, our investigation of potential slicing substrates was based on polyadenylated RNA-seq and small-RNA-seq and restricted to a high-confidence list of substrates based on previous publications. The application of degradome-sequencing to our samples might allow us to identify, with site-specific precision, new substrates that have not been previously reported, as well as substrates that may have been missed in our RNA-seq because they are not polyadenylated or do not increase significantly in *Ago2^CD/−^* tissues, owing to secondary changes or miRNA-mediated deadenylation.

## CONTACT FOR REAGENT AND RESOURCE SHARING

Further information and requests for resources and reagents should be directed to and will be fulfilled by the Lead Contact, Benjamin Kleaveland.

## MATERIALS AND METHODS

### Mouse husbandry

Mice were group-housed in a 12 hr light/dark cycle (light between 06:00 and 18:00) in a temperature-controlled room (21.1 ± 1.1°C) at Weill Cornell Medicine. Research with free access to water and food and maintained according to protocols approved by the Institutional Animal Care and Use Committee at Weill Cornell Medicine. Euthanasia of adult animals was performed by CO2 inhalation. Euthanasia of embryos was performed by decapitation. The developmental stage of embryos are indicated in the figure legends or methods. Sex was not determined for embryos or neonatal pups.

### Mouse strains and genotyping

The following mutant mouse strains were used in this study: *Ago2^CD/+^* [*Ago2tm3.1Ghan*/J] (JAX #014151);^19^ *Ago2^fl/fl^* [B6.129P2(129S4)-*Ago2tm1.1Tara*/J] (JAX #016520);^27^ *CMV-cre* [B6.Cg-Tg(*CMV-cre*)1Cgn/J] (JAX #006054);^73^ *VEC-cre* [B6.Cg-Tg(*Cdh5-cre*)1Spe/J] (JAX #033055);^74^ *Mlc2v-cre* [B6.129S4(Cg)-*Myl2tm1(cre)Krc*/AchakJ] (JAX #029465);^75^ *Actl6b-cre* [Tg(*Actl6b-Cre*)4092Jiwu/J] (JAX #027826);^76^ *ACTA1-cre* [B6.Cg-Tg(*ACTA1-cre*)79Jme/J] (JAX #006149);^63^ and *mTmG* [B6.129(Cg) - *Gt(ROSA)26Sortm4(ACTB-tdTomato,-EGFP)Luo*/J] (JAX #007676).^77^ The following inbred mouse strains were used in this study: DBA/2J (JAX #000671), 129X1/SvJ (JAX #000691), and C57BL/6J (JAX #000664). To generate the *Ago2^+/−^* mice used in this study, *Ago2^fl/fl^* mice harboring loxP sites flanking exons 9 and 11 were crossed to *CMV-cre* transgenic mice and then bred away from the *cre* transgene by mating to C57BL/6J animals for at least one generation. The recombined null allele lacks exons 9–11, and the resulting transcript contains a frame-shift coding sequence and a premature termination codon.

Genotyping was performed on genomic DNA extracted from mouse earsnips (adult mice), tails (neonates and embryos), or amniotic sac (embryos) using the HotSHOT method.^78^ PCR was performed with KAPA 2G Fast Genotyping Mix (Roche KK5121) and amplicons were analyzed by agarose gel electrophoresis. PCR primers, PCR conditions, and expected amplicon sizes are indicated in Table S10. Transnetyx MiniMUGA SNP analysis was also performed on select animals with *Ago2^CD^*, *Ago2^−^*, *Ago2^fl^* alleles to define the background strain; these mice were 96.3–98.5% C57BL6/J and C57BL/6NCrl (n = 5–6 backcrosses).

### Cell lines, cell culture, and live cell confocal microscopy

All cell lines were cultured at 37°C with 5% CO_2_. C2C12 cells are immortalized mouse myoblasts of female origin. C2C12 cells were cultured in DMEM, high glucose (Cytiva HyClone #SH30022) with 10% Tet-approved FBS (Gibco A7364) and 1X Pen/Strep (Gibco 15140-122) on tissue culture-treated plasticware. Cells were split every 1–3 days, maintaining confluency at less than 50–70% to minimize spontaneous differentiation and fusion. Polyclonal cell lines were generated as follows. Forward transfections were performed at 20% confluency with 2.6 μg of piggyBac plasmid DNA, 0.4 μg of Super piggyBac Transposase vector (System Biosciences PB210PA-1), and 7.5 μL of Lipofectamine 2000 (ThermoFisher 11668030), according to the manufacturer’s instructions. Transfection efficiencies were ∼20% based on control plasmids that constitutively expressed GFP or mCherry. Cells were cultured for 48 hours and then treated with 4 μg/mL puromycin (Gibco A1113803) for 48 hours to select for stably integrated expression cassettes. Polyclonal cell lines were cultured in the absence of puromycin for at least 72 hours before initiating any experiments. To induce expression of *Halo*, *Rtl1*, or *RTL1*, cell lines were treated with 1 μg/mL doxycycline (Fisher BP26531) for at least 24 hours. Doxycycline was replenished every 48 hours. For live cell confocal microscopy, polyclonal *2xFS-Halo* and *Rtl1-Halo* C2C12 cell lines were cultured in 8-well chamber slides (Ibidi), treated with doxycycline for 24 hours, stained with 200 nM membrane-permeable JF-646 Halo ligand (Promega HT1060) and 25 nM Hoechst (Apex Bio A3742) for ∼30 minutes at 37C, washed twice with fresh media, allowed to recover at 37C for 15 minutes, and then imaged by confocal microscopy (Zeiss LSM880) in an incubated chamber. For each image, 8–9 Z-slices were captured with a step-size of 1.01 microns. Z-stacks were projected based on average intensity. In the doxycycline-treated *2xFS-Halo* cell line, the voltages for the Green/GFP and Far-Red/JF-646 Halo ligand detectors were each reduced by ∼20% and attenuation values for the 488 nm and 633 nm excitation lasers were increased by ∼3% and ∼8%, respectively, to avoid saturating the image. Pinhole and scan speed parameters were identical for all images.

### Histology, immunohistochemistry, and immunofluorescence

Embryos and placentas were collected and fixed in 50 mL of 10% buffered formalin (VWR, 16004-121) for at least 24 hrs and stored in 70% ethanol. All staining, immunohistochemistry, and immunofluorescence was performed on formalin-fixed, paraffin-embedded 5–7 micron-thick tissue sections. Hematoxylin and eosin staining and Periodic Acid-Schiff staining (Abcam, ab-150680) were performed according to manufacturer’s instructions. Masson’s trichrome stains were performed on formalin-fixed, paraffin-embedded tissue sections. For Masson’s trichrome staining, sections were deparaffinized and fixed in Bouin’s solution overnight, stained with Weigert Iron Hematoxylin for 15 min, incubated in Biebrich Scarlet-Acid Fuchsin Stain for 5 min, incubated in Phosphomolybdic-Phosphotungstic Acid for 20 min, followed by staining with Aniline Blue for 2 min and acetic acid for 5–7 min. Slides were then dehydrated with ethanol and xylene.

Immunohistochemistry was performed on formalin-fixed, paraffin-embedded tissue sections using a Leica Bond automated staining system. Sections were pre-treated using heat-mediated antigen retrieval with sodium citrate buffer (pH 6.0) for 10 min. Slides were then incubated for 15 min at room temperature (RT) with anti-CD31/PECAM-1 antibody diluted 1:200 (Abcam, ab182981), followed by detection using an HRP-conjugated compact polymer system with 3,3′-Diaminobenzidine (DAB) used as a chromogen.

Immunofluorescence was performed on formalin-fixed, paraffin-embedded tissue sections as follows. Sections were pre-treated using heat-mediated antigen retrieval with sodium citrate buffer (pH 6.0) for 10 min. Sections were blocked with 2% normal donkey serum (Jackson ImmunoResearch 017-000-121) for one hour at RT, incubated overnight at 4C with anti-Endomucin antibody (clone V.7C7, Santa Cruz Biotechnology SC-65495) and anti-RTL1 antibody (YZ2844)(Ito 2015), each diluted 1:200 in PBST (PBS and 0.05% Tween-20), washed three times with PBST, and then incubated for one hour at RT with fluorophore-coupled donkey anti-rat IgG (Jackson ImmunoResearch 712-547-003) and donkey anti-rabbit IgG (Biotium 20152) secondary antibodies diluted 1:200 in PBST. Four images were collected per section using a Zeiss AxioObserver 7 Inverted Wide field/Fluorescence Microscope.

Histological images were visualized and analyzed using QuPath (v5.0). Placental capillary lumens and mouse muscle fiber cross-sectional areas were quantified by manually outlining the regions of interest (ROI) within each image. The area of each ROI was calculated using the software’s built-in measurement tools. To quantify placental capillary lumens, cross sections were measured in five 250,000 square micron tiles across the labyrinth zone. Muscle fiber cross-sectional areas were limited to 1–2 regions of intercostal muscle closest to the posterior of the animal.

### RNA extraction

Total RNA was extracted from flash frozen tissues collected from E15.5 and E19.5 embryos using TRI Reagent (ThermoFisher), according to manufacturer’s instructions with the following modifications. Mouse tissues were rapidly dissected after euthanasia and flash frozen in Eppendorf tubes in liquid N2. Tissue was transferred to a 15 mL conical tube, 1–2 mL of TRI Reagent was added, and the tissue was homogenized with a TissueRuptor (Qiagen) and disposable probes. Samples were transferred to Eppendorf tubes (1 mL per tube) and phase separated with 100 μL 1-bromo-3-chloropropane (J.T. Baker Analytical) per 1 mL of TRI Reagent. After isopropanol precipitation and two 70% ethanol washes, total RNA was resuspended in RNase-free water.

### RT–qPCR

For mouse tissues, 0.5–1 μg total RNA was treated with dsDNase, then reverse transcribed with Maxima H Minus First Strand cDNA Synthesis Kit (ThermoFisher K1682) with a mix of oligo(dT) primers and random hexamers following manufacturer’s instructions. Expression of *Rtl1* was measured by qPCR using PowerUP SYBR Green Master Mix (Applied Biosystems A25742) on a QuantStudio5 system (Applied Biosystems) and quantified using the ΔΔCT method with the geometric mean of *Acta1*, *Actb*, and *Gapdh* expression as the internal normalization control. qPCR primers are listed in Table S10.

### RNA-Seq and analysis

For all samples (n = 3–4 biological replicates per genotype, with the exception of E19.5 *Ago2^+/+^* placenta and E19.5 *Acta1-cre*; *Ago2^+/fl^* diaphragm for which n = 2), the NEBNext Ultra II Directional RNA Library Prep Kit (NEB E7760) was used to generate stranded, poly(A)-selected RNA-seq libraries. Briefly, 1 μg of total RNA was poly(A) enriched and then reverse transcribed into double-strand cDNA. The cDNA samples were fragmented, end-repaired, and polyadenylated before ligation of TruSeq adapters containing an index sequence for multiplex sequencing. Multiplexing fragments containing TruSeq adapters on both ends were selectively enriched with 8–9 PCR cycles. All libraries were sequenced on the NovaseqX (Illumina) with 100 nt paired-end reads and ∼20 million reads per sample.

Alignment and differential expression of the RNA-seq libraries was performed using the nf-core pipeline.^79,80^ Briefly, the default RNAseq pipeline was used, specifying STAR alignment and using a salmon counter. Reads were aligned to the mouse genome (mm39) using STAR v2.7.1a^81^ and Gencode annotations (m39.gencode.vM25.basic.annotation.gtf; downloaded 7/10/23). For mouse tissues, the gtf file was modified to include *Rtl1as* using the same coordinates as *Rtl1* exon 2. For C2C12 cell lines, the mouse genome and gtf annotation file were modified to include the corresponding piggyBac sequences and either *Rtl1*, *RTL1*, or *Halo* transgenes as an artificial chromosome. As no reads mapped to the endogenous *Rtl1* locus in *Halo*-expressing C2C12 cells, we excluded endogenous *Rtl1* from any downstream analysis.

Count files were merged to generate tables of gene counts for each tissue organized by genotype. For mouse tissue libraries, salmon.merged.gene_counts_length_scaled.tsv were imported directly into DESeq2; for C2C12 libraries, salmon.merged.gene_counts.tsv and salmon.merged.gene_lengths.tsv were imported into DESeq2 via tximport. Differential expression was performed using DESeq2 without the lfcShrink function.^82^ RNA-seq browser tracks were visualized in the UCSC genome browser and IGV v2.3.34.^83,84^ TPMs provided in Table S4 were based on salmon.merged.gene_tpm.tsv files. For RNA-seq MA plots, only RNAs with at least 50 mean normalized counts in the control samples are plotted.

Principal component analysis was performed on DESeq2 normalized counts from each sample using the prcomp function in R, with genes filtered to include those with the highest variance across samples (500 most variable genes). The first two principal components represent the largest sources of variation in the dataset and were used for visualization.

Gene set enrichment analysis (GSEA) was performed using WebGestalt^85^ and gene ontology functional databases for cell components and molecular function with the following parameters: minimum number of IDs in category, 20; maximum number of IDs in category, 1000; significance level, FDR < 0.05; permutations, 1000. The input for each GSEA was a list of *M. musculus* gene IDs ranked by STAT, which is the log2 fold-change divided by log2 fold-change standard error. Overrepresentation analysis was performed using Enrichr^64,65^ with the differentially expressed upregulated or downregulated gene sets from each comparison, a background gene set consisting all genes detected in the corresponding comparison, and the 2022 ChIP-X Enrichment Analysis database containing 757 gene sets covering >18,000 genes. Only gene sets from mouse cell lines or tissues with at least five genes overlapping our input gene set were plotted.

### miRNA target analysis

miRNA target predictions were downloaded from TargetScanMouse Release 8.0.^32^ Differential expression data (DESeq2 output), filtered to exclude genes with a mean normalized count < 50, were analyzed for repression of genes predicted to be miRNA targets. For each miRNA, three sets of predicted targets were analyzed: all predicted targets, conserved predicted targets (unless the miRNA wasn’t conserved), and top predicted targets. Top predicted targets were defined as the top 10% of all predicted targets based on cumulative weighted context++ scores.^31^ Each of the three sets of targets was compared to nontargets. The nontarget set was selected as follows. First, 3′ UTR sequences from TargetScanMouse Release 8.0 (downloaded on 7/10/24) were used to group genes into 10 bins based on 3′ UTR length. Next, for each target, one transcript not predicted to be a target of the miRNA family was selected, with replacement, from the corresponding UTR bin. For each target set, the distribution of log2 fold-changes in experimental samples relative to control samples, as determined by DESeq2, was compared to the distribution of log2 fold-changes of the 3′ UTR length-matched nontarget set, and statistical significance was determined using a Mann-Whitney test. The degree of repression is represented by subtracting the median log2 fold-change of the target set from the median log2 fold-change of its 3′ UTR length-matched nontarget set. The analysis described above was repeated 20 additional times for each target set, and the mean difference in median log2 fold-changes and the median *P* value across the 21 replicates were plotted. For simplicity, only the nontarget set with the median *P* value in the all targets analysis was shown in each cumulative distribution function plot.

### Small-RNA-seq and analysis

Small-RNA sequencing libraries were prepared with 2 μg of total RNA from each diaphragm (n = 2–3 biological replicates per genotype). RNAs that co-migrated within the range of 17 and 32 nt radiolabeled internal standards run in adjacent lanes were isolated on a 15% polyacrylamide urea gel. Gel slices were macerated and RNA was eluted overnight at 4C in 400 μL of 0.3 M NaCl. After Spin-X column (Corning CLS8170) cleanup, RNA was precipitated for >2 hours at – 20C with 3 μL of GlycoBlue (ThermoFisher AM9515) and 1 mL of 100% ethanol and then resuspended in 8 μL RNase-free H2O. All 8 μL were used as input for the NEBNext Low-bias

Small-RNA Library Prep Kit (NEB E3420). Libraries were prepared, according to manufacturer’s instructions, with 1:3 dilutions of both 5′ and 3′ adapters. After amplifying libraries with 11 cycles of PCR, bead-based size selection was performed to further enrich libraries with miRNA-sized inserts. Libraries were sequenced on a NovaSeqX (Illumina) with single-end 50 bp reads and ∼26 million reads per sample.

Reads were trimmed of adaptor sequence using cutadapt and filtered using fastq_quality_filter (FastX Toolkit; http://hannonlab.cshl.edu/fastx_toolkit/) with the parameters “-q 30 –p 100” to ensure that all bases had an accuracy of 99.9%. The first 19 nt of each read were matched to a dictionary of miRNA sequences downloaded from TargetScanMouse Release 8.0, requiring no mismatches between the read and the miRNA dictionary, using custom python scripts.

Differential expression of the small-RNA-seq libraries was performed using the nf-core 24.10.1 pipeline and DESeq2 without the lfcShrink function and independent filtering = false. For small-RNA-seq MA plots, only RNAs with at least 5 mean normalized counts in the control samples are plotted.

### Small RNA northern blots

For each E15.5 liver sample, 10 μg of total RNA was resolved on a 15% polyacrylamide urea gel, transferred to a Hybond-NX membrane (GE Healthcare) using a semi-dry transfer apparatus (Bio-Rad), and incubated at 60C for 1 hour with EDC (1-(3-Dimethylaminopropyl)-3-ethylcarbodiimide; TCI 1892-57-5) diluted in 1X methylimidazole to chemically crosslink 5′ phosphates to the membrane. Blots were blocked in ULTRAhyb-Oligo (ThermoFisher AM8663) for at least 10 min, hybridized overnight with radiolabeled DNA probes (Table S10) diluted in ULTRAhyb-Oligo, and then washed 3×20 min with low-stringency buffer (2X SSC, 0.1% SDS) and 1×10 min with high-stringency buffer (0.1X SSC, 0.1% SDS). All steps were performed in a hybridization oven set to 42C. Northern blots were exposed to a Phosphorimager plate for at least 24 hours, imaged using a Typhoon Phosphorimager (Cytiva) and analyzed with ImageQuant TL (v8.1.0.0). Blots were stripped by incubating 4×20 min in boiling-hot 0.04% SDS, reimaged, and then reprobed.

### Protein extraction and western blots

Mouse livers (n = 2–3 per genotype) were dissected from E15.5 embryos and flash frozen in liquid N2. Frozen tissues were lysed in 500 μL 1X RIPA buffer supplemented with 2 μL of Protease Inhibitor cocktail III (MilliporeSigma 539134) and 5 μL of Turbo DNase (ThermoFisher AM2238), probe sonicated for 2 min over ice until fully lysed, and then clarified by centrifugation at 15,000 x g for 5 min. For C2C12 cell lines, cells were trypsinized 24 hours after doxycycline induction, pelleted by centrifugation, and washed with 1X PBS. The cell pellets were mixed with 500 μL of RIPA buffer supplemented with 2 μL of Protease Inhibitor Cocktail III and 5 μL of Turbo DNAse, lysed for 10 min over ice, and then clarified by centrifugation at 15,000 x g for 5 min. Total protein was quantified using the Pierce BCA Protein Assay (ThermoFisher 23225).

For each western blot, samples were diluted with RIPA buffer to equal volumes, then mixed with 1X NuPAGE LDS Sample Buffer (ThermoFisher NP0008) and 50 mM DTT. Samples were heated for 5 min at 95C, loaded on a NuPAGE Tris-Glycine 4–20% protein gel (ThermoFisher XP04205BOX) in 1X Tris-Glycine, 10% SDS Running Buffer, and run at 190V for approximately 45 min. Using Tris-Glycine supplemented with 10% methanol, the gel was subsequently transferred to a nitrocellulose membrane (Cytiva 10600003) at 30V overnight at 4C.

The membrane was blocked for one hour with Intercept (TBS) Protein-Free Blocking Buffer (LICORbio 927-80003) then incubated with primary antibody overnight at 4°C on a shaker. The next day, the membrane was washed 3x with 1X TBST, and then incubated with secondary antibody for one hour at room temperature. The membrane was washed 3x with 1X TBST and 1x with 1X TBS, and then imaged in the 800 channel (Bio-Rad ChemiDoc MP Imaging System). Afterwards, the membrane was stained for total protein using the Revert 700 Total Protein Stain (LICORbio 926-11016) and imaged in the 700 channel. The primary antibodies used in this study were anti-AGO2 (1:1000, ThermoFisher MA5-32520), anti-RTL1 (1:1000, YZ2844),^55^ and anti-alpha Tubulin (1:1000, Abcam ab52866). The secondary antibody used in this study was IRDye 800CW Goat anti-Rabbit (1:10000, LI-CORE 926032211). AGO2 band intensity and total protein intensity were measured on the raw TIF files in ImageJ, first using local background subtraction, and then normalized to the average WT intensity.

## QUANTIFICATION AND STATISTICAL ANALYSIS

Graphs were generated in GraphPad Prism 11, R, and statistical analyses were performed using GraphPad Prism 11, Excel, or R. Statistical parameters including the value of n, statistical test, and statistical significance (*P* value) are reported in the figures or their legends. For studies involving mouse tissues, replicates refer to samples derived from different mice. No statistical methods were used to predetermine sample size. Statistical tests were selected based on the desired comparison. Unpaired two-tailed t-tests without correction for multiple-hypothesis testing were used to assess significance when comparing the frequency of mean cross-sectional areas and vessel lumenal areas in each bin, and when comparing TPM levels between two groups.

One-way ANOVA was used to assess significance when comparing measurements across four genotypes (e.g. RT-qPCR, central nuclei, etc); significant ANOVA results were followed by appropriate post-hoc testing and correction for multiple-hypothesis testing, as determined by the data type. For differential expression of global measurements (RNA-seq, small-RNA-seq), we report DESeq2-generated adjusted *P* values using the Wald test with the Benjamini-Hochberg correction for multiple-hypothesis testing. The Mann-Whitney test was used to compare cumulative distributions of mRNA fold-changes between two gene sets.

## DATA AND SOFTWARE AVAILABILITY

Sequencing datasets generated in this study have been deposited in the GEO under accession numbers (small-RNA-seq) and (RNA-seq).

## ACKNOWLEDGMENTS

We thank E. Lai, H. Stuhlman, C. Lima, M. Baylies and members of the Kleaveland lab for helpful discussions; A. DiLorenzo for providing *VEC-cre* and *Mlc2v-cre* mice; C. Stewart for providing anti-mouse RTL1 antibody; M. Baylies for providing C2C12 myoblasts; the Center for Translational Pathology and Histology Core for histology services and consultation; the WCM Genomics Resource Core Facility for sequencing; the WCM Image Analysis and Microscopy Core for microscopy expertise; the Research Animal Resource Center for animal husbandry.

This work was supported by National Institutes of Health grants R35GM147463 (B.K.) and R35GM147463-S1 (N.A.) from the National Institute of General Medical Sciences. The content is solely the responsibility of the authors and does not necessarily represent the official views of the National Institutes of Health.

## AUTHOR CONTRIBUTIONS

N.A. and B.K. conceived the project and designed the study. N.A. collected embryos/tissues for RNA and histology, prepared and analyzed RNA-seq libraries from mouse tissues, analyzed small-RNA-seq libraries, performed immunofluorescence and some histology stains, and performed all tissue morphometric analyses. C.D. performed western blotting and analyzed RNA-seq libraries from C2C12 cells. B.K. performed northern blots, generated polyclonal C2C12 cell lines, and prepared RNA-seq libraries from C2C12 cells and small-RNA-seq libraries. N.A. and B.K. collected mouse survival data. N.A. and B.K. prepared the figures. B.K. drafted the manuscript, and N.A. and B.K. revised the manuscript.

## REFERENCES

1. Shabalina, S.A., and Koonin, E.V. (2008). Origins and evolution of eukaryotic RNA interference. Trends Ecol Evol 23, 578–587. 10.1016/j.tree.2008.06.005.

2. Makarova, K.S., Wolf, Y.I., van der Oost, J., and Koonin, E.V. (2009). Prokaryotic homologs of Argonaute proteins are predicted to function as key components of a novel system of defense against mobile genetic elements. Biol Direct 4, 29. 10.1186/1745-6150-4-29.

3. Obbard, D.J., Gordon, K.H., Buck, A.H., and Jiggins, F.M. (2009). The evolution of RNAi as a defence against viruses and transposable elements. Philos Trans R Soc Lond B Biol Sci 364, 99–115. 10.1098/rstb.2008.0168.

4. Olovnikov, I.A., and Kalmykova, A.I. (2013). piRNA clusters as a main source of small RNAs in the animal germline. Biochemistry (Mosc) 78, 572–584. 10.1134/S0006297913060035.

5. Cerutti, H., and Casas-Mollano, J.A. (2006). On the origin and functions of RNA-mediated silencing: from protists to man. Curr Genet 50, 81–99. 10.1007/s00294-006-0078-x.

6. Swarts, D.C., Makarova, K., Wang, Y., Nakanishi, K., Ketting, R.F., Koonin, E.V., Patel, D.J., and van der Oost, J. (2014). The evolutionary journey of Argonaute proteins. Nat Struct Mol Biol 21, 743–753. 10.1038/nsmb.2879.

7. Bartel, D.P. (2018). Metazoan MicroRNAs. Cell 173, 20–51. 10.1016/j.cell.2018.03.006.

8. Pijuan-Sala, B., Wilson, N.K., Xia, J., Hou, X., Hannah, R.L., Kinston, S., Calero-Nieto, F.J., Poirion, O., Preissl, S., Liu, F., and Gottgens, B. (2020). Single-cell chromatin accessibility maps reveal regulatory programs driving early mouse organogenesis. Nat Cell Biol 22, 487–497. 10.1038/s41556-020-0489-9.

9. Liu, J., Carmell, M.A., Rivas, F.V., Marsden, C.G., Thomson, J.M., Song, J.J., Hammond, S.M., Joshua-Tor, L., and Hannon, G.J. (2004). Argonaute2 is the catalytic engine of mammalian RNAi. Science 305, 1437–1441. 10.1126/science.1102513.

10. Meister, G., Landthaler, M., Patkaniowska, A., Dorsett, Y., Teng, G., and Tuschl, T. (2004). Human Argonaute2 mediates RNA cleavage targeted by miRNAs and siRNAs. Mol Cell 15, 185–197. 10.1016/j.molcel.2004.07.007.

11. Rand, T.A., Ginalski, K., Grishin, N.V., and Wang, X. (2004). Biochemical identification of Argonaute 2 as the sole protein required for RNA-induced silencing complex activity. Proc Natl Acad Sci U S A 101, 14385–14389. 10.1073/pnas.0405913101.

12. Park, M.S., Sim, G., Kehling, A.C., and Nakanishi, K. (2020). Human Argonaute2 and Argonaute3 are catalytically activated by different lengths of guide RNA. Proc Natl Acad Sci U S A 117, 28576–28578. 10.1073/pnas.2015026117.

13. Zhang, H., Sim, G., Kehling, A.C., Adhav, V.A., Savidge, A., Pastore, B., Tang, W., and Nakanishi, K. (2024). Target cleavage and gene silencing by Argonautes with cityRNAs. Cell Rep 43, 114806. 10.1016/j.celrep.2024.114806.

14. Hammond, S.M., Bernstein, E., Beach, D., and Hannon, G.J. (2000). An RNA-directed nuclease mediates post-transcriptional gene silencing in Drosophila cells. Nature 404, 293–296. 10.1038/35005107.

15. Zamore, P.D., Tuschl, T., Sharp, P.A., and Bartel, D.P. (2000). RNAi: double-stranded RNA directs the ATP-dependent cleavage of mRNA at 21 to 23 nucleotide intervals. Cell 101, 25–33. 10.1016/S0092-8674(00)80620-0.

16. Elbashir, S.M., Harborth, J., Lendeckel, W., Yalcin, A., Weber, K., and Tuschl, T. (2001). Duplexes of 21-nucleotide RNAs mediate RNA interference in cultured mammalian cells. Nature 411, 494–498. 10.1038/35078107.

17. Adams, D., Gonzalez-Duarte, A., O’Riordan, W.D., Yang, C.C., Ueda, M., Kristen, A.V., Tournev, I., Schmidt, H.H., Coelho, T., Berk, J.L., et al. (2018). Patisiran, an RNAi Therapeutic, for Hereditary Transthyretin Amyloidosis. N Engl J Med 379, 11–21. 10.1056/NEJMoa1716153.

18. Jadhav, V., Vaishnaw, A., Fitzgerald, K., and Maier, M.A. (2024). RNA interference in the era of nucleic acid therapeutics. Nat Biotechnol 42, 394–405. 10.1038/s41587-023-02105-y.

19. Cheloufi, S., Dos Santos, C.O., Chong, M.M., and Hannon, G.J. (2010). A dicer-independent miRNA biogenesis pathway that requires Ago catalysis. Nature 465, 584–589. 10.1038/nature09092.

20. Jee, D., Yang, J.S., Park, S.M., Farmer, D.T., Wen, J., Chou, T., Chow, A., McManus, M.T., Kharas, M.G., and Lai, E.C. (2018). Dual Strategies for Argonaute2-Mediated Biogenesis of Erythroid miRNAs Underlie Conserved Requirements for Slicing in Mammals. Mol Cell 69, 265–278 e266. 10.1016/j.molcel.2017.12.027.

21. Kumar, M., Maria, A.G., Prajapat, M., and Vidigal, J.A. (2025). AGO2 slicing of a domesticated retrotransposon is necessary for normal vasculature development. bioRxiv. 10.1101/2025.04.02.646793.

22. Cifuentes, D., Xue, H., Taylor, D.W., Patnode, H., Mishima, Y., Cheloufi, S., Ma, E., Mane, S., Hannon, G.J., Lawson, N.D., et al. (2010). A novel miRNA processing pathway independent of Dicer requires Argonaute2 catalytic activity. Science 328, 1694–1698. 10.1126/science.1190809.

23. Yang, J.S., Maurin, T., Robine, N., Rasmussen, K.D., Jeffrey, K.L., Chandwani, R., Papapetrou, E.P., Sadelain, M., O’Carroll, D., and Lai, E.C. (2010). Conserved vertebrate mir-451 provides a platform for Dicer-independent, Ago2-mediated microRNA biogenesis. Proc Natl Acad Sci U S A 107, 15163–15168. 10.1073/pnas.1006432107.

24. Seitz, H., Youngson, N., Lin, S.P., Dalbert, S., Paulsen, M., Bachellerie, J.P., Ferguson-Smith, A.C., and Cavaille, J. (2003). Imprinted microRNA genes transcribed antisense to a reciprocally imprinted retrotransposon-like gene. Nat Genet 34, 261–262. 10.1038/ng1171.

25. Davis, E., Caiment, F., Tordoir, X., Cavaille, J., Ferguson-Smith, A., Cockett, N., Georges, M., and Charlier, C. (2005). RNAi-mediated allelic trans-interaction at the imprinted Rtl1/Peg11 locus. Curr Biol 15, 743–749. 10.1016/j.cub.2005.02.060.

26. Sekita, Y., Wagatsuma, H., Nakamura, K., Ono, R., Kagami, M., Wakisaka, N., Hino, T., Suzuki-Migishima, R., Kohda, T., Ogura, A., et al. (2008). Role of retrotransposon-derived imprinted gene, Rtl1, in the feto-maternal interface of mouse placenta. Nat Genet 40, 243–248. 10.1038/ng.2007.51.

27. O’Carroll, D., Mecklenbrauker, I., Das, P.P., Santana, A., Koenig, U., Enright, A.J., Miska, E.A., and Tarakhovsky, A. (2007). A Slicer-independent role for Argonaute 2 in hematopoiesis and the microRNA pathway. Genes Dev 21, 1999–2004. 10.1101/gad.1565607.

28. Rasmussen, K.D., Simmini, S., Abreu-Goodger, C., Bartonicek, N., Di Giacomo, M., Bilbao-Cortes, D., Horos, R., Von Lindern, M., Enright, A.J., and O’Carroll, D. (2010). The miR-144/451 locus is required for erythroid homeostasis. J Exp Med 207, 1351–1358. 10.1084/jem.20100458.

29. Patrick, D.M., Zhang, C.C., Tao, Y., Yao, H., Qi, X., Schwartz, R.J., Jun-Shen Huang, L., and Olson, E.N. (2010). Defective erythroid differentiation in miR-451 mutant mice mediated by 14-3-3zeta. Genes Dev 24, 1614–1619. 10.1101/gad.1942810.

30. Xu, P., Palmer, L.E., Lechauve, C., Zhao, G., Yao, Y., Luan, J., Vourekas, A., Tan, H., Peng, J., Schuetz, J.D., et al. (2019). Regulation of gene expression by miR-144/451 during mouse erythropoiesis. Blood 133, 2518–2528. 10.1182/blood.2018854604.

31. Agarwal, V., Bell, G.W., Nam, J.W., and Bartel, D.P. (2015). Predicting effective microRNA target sites in mammalian mRNAs. Elife 4. 10.7554/eLife.05005.

32. McGeary, S.E., Lin, K.S., Shi, C.Y., Pham, T.M., Bisaria, N., Kelley, G.M., and Bartel, D.P. (2019). The biochemical basis of microRNA targeting efficacy. Science 366. 10.1126/science.aav1741.

33. Wang, D., Zhang, Z., O’Loughlin, E., Lee, T., Houel, S., O’Carroll, D., Tarakhovsky, A., Ahn, N.G., and Yi, R. (2012). Quantitative functions of Argonaute proteins in mammalian development. Genes Dev 26, 693–704. 10.1101/gad.182758.111.

34. Johnson, K.C., and Corey, D.R. (2023). RNAi in cell nuclei: potential for a new layer of biological regulation and a new strategy for therapeutic discovery. RNA 29, 415–422. 10.1261/rna.079500.122.

35. Turgeon, B., and Meloche, S. (2009). Interpreting neonatal lethal phenotypes in mouse mutants: insights into gene function and human diseases. Physiol Rev 89, 1–26. 10.1152/physrev.00040.2007.

36. Perez-Garcia, V., Fineberg, E., Wilson, R., Murray, A., Mazzeo, C.I., Tudor, C., Sienerth, A., White, J.K., Tuck, E., Ryder, E.J., et al. (2018). Placentation defects are highly prevalent in embryonic lethal mouse mutants. Nature 555, 463–468. 10.1038/nature26002.

37. Yekta, S., Shih, I.H., and Bartel, D.P. (2004). MicroRNA-directed cleavage of HOXB8 mRNA. Science 304, 594–596. 10.1126/science.1097434.

38. Shin, C., Nam, J.W., Farh, K.K., Chiang, H.R., Shkumatava, A., and Bartel, D.P. (2010). Expanding the microRNA targeting code: functional sites with centered pairing. Mol Cell 38, 789–802. 10.1016/j.molcel.2010.06.005.

39. Karginov, F.V., Cheloufi, S., Chong, M.M., Stark, A., Smith, A.D., and Hannon, G.J. (2010). Diverse endonucleolytic cleavage sites in the mammalian transcriptome depend upon microRNAs, Drosha, and additional nucleases. Mol Cell 38, 781–788. 10.1016/j.molcel.2010.06.001.

40. Bracken, C.P., Szubert, J.M., Mercer, T.R., Dinger, M.E., Thomson, D.W., Mattick, J.S., Michael, M.Z., and Goodall, G.J. (2011). Global analysis of the mammalian RNA degradome reveals widespread miRNA-dependent and miRNA-independent endonucleolytic cleavage. Nucleic Acids Res 39, 5658–5668. 10.1093/nar/gkr110.

41. Hansen, T.B., Wiklund, E.D., Bramsen, J.B., Villadsen, S.B., Statham, A.L., Clark, S.J., and Kjems, J. (2011). miRNA-dependent gene silencing involving Ago2-mediated cleavage of a circular antisense RNA. EMBO J 30, 4414–4422. 10.1038/emboj.2011.359.

42. Kleaveland, B., Shi, C.Y., Stefano, J., and Bartel, D.P. (2018). A Network of Noncoding Regulatory RNAs Acts in the Mammalian Brain. Cell 174, 350–362 e317. 10.1016/j.cell.2018.05.022.

43. Rishik, S., Hirsch, P., Grandke, F., Fehlmann, T., and Keller, A. (2025). miRNATissueAtlas 2025: an update to the uniformly processed and annotated human and mouse non-coding RNA tissue atlas. Nucleic Acids Res 53, D129–D137. 10.1093/nar/gkae1036.

44. Alexander, M.S., Casar, J.C., Motohashi, N., Myers, J.A., Eisenberg, I., Gonzalez, R.T., Estrella, E.A., Kang, P.B., Kawahara, G., and Kunkel, L.M. (2011). Regulation of DMD pathology by an ankyrin-encoded miRNA. Skelet Muscle 1, 27. 10.1186/2044-5040-1-27.

45. Alexander, M.S., Casar, J.C., Motohashi, N., Vieira, N.M., Eisenberg, I., Marshall, J.L., Gasperini, M.J., Lek, A., Myers, J.A., Estrella, E.A., et al. (2014). MicroRNA-486-dependent modulation of DOCK3/PTEN/AKT signaling pathways improves muscular dystrophy-associated symptoms. J Clin Invest 124, 2651–2667. 10.1172/JCI73579.

46. Samani, A., Hightower, R.M., Reid, A.L., English, K.G., Lopez, M.A., Doyle, J.S., Conklin, M.J., Schneider, D.A., Bamman, M.M., Widrick, J.J., et al. (2022). miR-486 is essential for muscle function and suppresses a dystrophic transcriptome. Life Sci Alliance 5. 10.26508/lsa.202101215.

47. Tam, O.H., Aravin, A.A., Stein, P., Girard, A., Murchison, E.P., Cheloufi, S., Hodges, E., Anger, M., Sachidanandam, R., Schultz, R.M., and Hannon, G.J. (2008). Pseudogene-derived small interfering RNAs regulate gene expression in mouse oocytes. Nature 453, 534–538. 10.1038/nature06904.

48. Watanabe, T., Totoki, Y., Toyoda, A., Kaneda, M., Kuramochi-Miyagawa, S., Obata, Y., Chiba, H., Kohara, Y., Kono, T., Nakano, T., et al. (2008). Endogenous siRNAs from naturally formed dsRNAs regulate transcripts in mouse oocytes. Nature 453, 539–543. 10.1038/nature06908.

49. Suh, N., Baehner, L., Moltzahn, F., Melton, C., Shenoy, A., Chen, J., and Blelloch, R. (2010). MicroRNA function is globally suppressed in mouse oocytes and early embryos. Curr Biol 20, 271–277. 10.1016/j.cub.2009.12.044.

50. Ma, J., Flemr, M., Stein, P., Berninger, P., Malik, R., Zavolan, M., Svoboda, P., and Schultz, R.M. (2010). MicroRNA activity is suppressed in mouse oocytes. Curr Biol 20, 265–270. 10.1016/j.cub.2009.12.042.

51. Flemr, M., Malik, R., Franke, V., Nejepinska, J., Sedlacek, R., Vlahovicek, K., and Svoboda, P. (2013). A retrotransposon-driven dicer isoform directs endogenous small interfering RNA production in mouse oocytes. Cell 155, 807–816. 10.1016/j.cell.2013.10.001.

52. Stein, P., Rozhkov, N.V., Li, F., Cardenas, F.L., Davydenko, O., Vandivier, L.E., Gregory, B.D., Hannon, G.J., and Schultz, R.M. (2015). Essential Role for endogenous siRNAs during meiosis in mouse oocytes. PLoS Genet 11, e1005013. 10.1371/journal.pgen.1005013.

53. Sala, L., Kumar, M., Prajapat, M., Chandrasekhar, S., Cosby, R.L., La Rocca, G., Macfarlan, T.S., Awasthi, P., Chari, R., Kruhlak, M., and Vidigal, J.A. (2023). AGO2 silences mobile transposons in the nucleus of quiescent cells. Nat Struct Mol Biol 30, 1985–1995. 10.1038/s41594-023-01151-z.

54. Tam, O.H., Forcier, T., Gale Hammell, M. (2026). Custom GTF generation for TEToolkit (Version 1.0). 10.5281/zenodo.18927862.

55. Ito, M., Sferruzzi-Perri, A.N., Edwards, C.A., Adalsteinsson, B.T., Allen, S.E., Loo, T.H., Kitazawa, M., Kaneko-Ishino, T., Ishino, F., Stewart, C.L., and Ferguson-Smith, A.C. (2015). A trans-homologue interaction between reciprocally imprinted miR-127 and Rtl1 regulates placenta development. Development 142, 2425–2430. 10.1242/dev.121996.

56. Kitazawa, M., Hayashi, S., Imamura, M., Takeda, S., Oishi, Y., Kaneko-Ishino, T., and Ishino, F. (2020). Deficiency and overexpression of Rtl1 in the mouse cause distinct muscle abnormalities related to Temple and Kagami-Ogata syndromes. Development 147. 10.1242/dev.185918.

57. Kagami, M., Yamazawa, K., Matsubara, K., Matsuo, N., and Ogata, T. (2008). Placentomegaly in paternal uniparental disomy for human chromosome 14. Placenta 29, 760–761. 10.1016/j.placenta.2008.06.001.

58. Ogata, T., and Kagami, M. (2016). Kagami-Ogata syndrome: a clinically recognizable upd(14)pat and related disorder affecting the chromosome 14q32.2 imprinted region. J Hum Genet 61, 87–94. 10.1038/jhg.2015.113.

59. Prasasya, R., Grotheer, K.V., Siracusa, L.D., and Bartolomei, M.S. (2020). Temple syndrome and Kagami-Ogata syndrome: clinical presentations, genotypes, models and mechanisms. Hum Mol Genet 29, R107–R116. 10.1093/hmg/ddaa133.

60. Fu, Y., Zeng, X., Liu, Y., Jia, S., Jiang, Y., Tan, J.P., Yuan, Y., Xia, T., Mei, Y., Wen, S., et al. (2026). A spatiotemporal transcriptomic atlas of the mouse placenta reveals glycogen cell-mediated metabolic support essential for fetal viability. eLife Sciences Publications, Ltd.

61. Sequeira, C., Wackerbarth, L.M., Pena, A., Sa-Pereira, M., Franco, C.A., and Gomes, E.R. (2024). Myonuclear position and blood vessel organization during skeletal muscle postnatal development. Development 151. 10.1242/dev.202548.

62. Malicdan, M.C., and Nishino, I. (2012). Autophagy in lysosomal myopathies. Brain Pathol 22, 82–88. 10.1111/j.1750-3639.2011.00543.x.

63. Miniou, P., Tiziano, D., Frugier, T., Roblot, N., Le Meur, M., and Melki, J. (1999). Gene targeting restricted to mouse striated muscle lineage. Nucleic Acids Res 27, e27. 10.1093/nar/27.19.e27.

64. Keenan, A.B., Torre, D., Lachmann, A., Leong, A.K., Wojciechowicz, M.L., Utti, V., Jagodnik, K.M., Kropiwnicki, E., Wang, Z., and Ma’ayan, A. (2019). ChEA3: transcription factor enrichment analysis by orthogonal omics integration. Nucleic Acids Res 47, W212–w224. 10.1093/nar/gkz446.

65. Li, L., Sheng, P., Li, T., Fields, C.J., Hiers, N.M., Wang, Y., Li, J., Guardia, C.M., Licht, J.D., and Xie, M. (2021). Widespread microRNA degradation elements in target mRNAs can assist the encoded proteins. Genes Dev 35, 1595–1609. 10.1101/gad.348874.121.

66. Song, X., Niu, L., Fan, X., Xu, X., Zhao, Z., Zhang, Z., Tong, Y., Huang, H., Zhu, Z., Cheng, H., et al. (2026). Overexpression of Rtl1 via synonymous mutation silencing miRNA target site drives skeletal muscle hypertrophy and inflammation in mice. BMC Biol 24. 10.1186/s12915-026-02534-6.

67. Ogasawara, M., and Nishino, I. (2023). A review of major causative genes in congenital myopathies. J Hum Genet 68, 215–225. 10.1038/s10038-022-01045-w.

68. Jungbluth, H., Treves, S., Zorzato, F., Sarkozy, A., Ochala, J., Sewry, C., Phadke, R., Gautel, M., and Muntoni, F. (2018). Congenital myopathies: disorders of excitation-contraction coupling and muscle contraction. Nat Rev Neurol 14, 151–167. 10.1038/nrneurol.2017.191.

69. Segel, M., Lash, B., Song, J., Ladha, A., Liu, C.C., Jin, X., Mekhedov, S.L., Macrae, R.K., Koonin, E.V., and Zhang, F. (2021). Mammalian retrovirus-like protein PEG10 packages its own mRNA and can be pseudotyped for mRNA delivery. Science 373, 882–889. 10.1126/science.abg6155.

70. Kitazawa, M., Tamura, M., Kaneko-Ishino, T., and Ishino, F. (2017). Severe damage to the placental fetal capillary network causes mid- to late fetal lethality and reduction in placental size in Peg11/Rtl1 KO mice. Genes Cells 22, 174–188. 10.1111/gtc.12465.

71. Mullokandov, G., Baccarini, A., Ruzo, A., Jayaprakash, A.D., Tung, N., Israelow, B., Evans, M.J., Sachidanandam, R., and Brown, B.D. (2012). High-throughput assessment of microRNA activity and function using microRNA sensor and decoy libraries. Nat Methods 9, 840–846. 10.1038/nmeth.2078.

72. Kraus, C., Wang, J., Zheng, H., Broderick, J., Ajaykumar, N., Zamani, M., Yang, M., Cecchini, K., Liang, S.Q., Kolumba, O., et al. (2025). Absolute quantification of mammalian microRNAs for therapeutic RNA cleavage and detargeting. RNA 31, 1081–1090. 10.1261/rna.080566.125.

73. Schwenk, F., Baron, U., and Rajewsky, K. (1995). A cre-transgenic mouse strain for the ubiquitous deletion of loxP-flanked gene segments including deletion in germ cells. Nucleic Acids Res 23, 5080–5081. 10.1093/nar/23.24.5080.

74. Chen, M.J., Yokomizo, T., Zeigler, B.M., Dzierzak, E., and Speck, N.A. (2009). Runx1 is required for the endothelial to haematopoietic cell transition but not thereafter. Nature 457, 887–891. 10.1038/nature07619.

75. Chen, J., Kubalak, S.W., Minamisawa, S., Price, R.L., Becker, K.D., Hickey, R., Ross, J., Jr., and Chien, K.R. (1998). Selective requirement of myosin light chain 2v in embryonic heart function. J Biol Chem 273, 1252–1256. 10.1074/jbc.273.2.1252.

76. Zhan, X., Cao, M., Yoo, A.S., Zhang, Z., Chen, L., Crabtree, G.R., and Wu, J.I. (2015). Generation of BAF53b-Cre transgenic mice with pan-neuronal Cre activities. Genesis 53, 440–448. 10.1002/dvg.22866.

77. Muzumdar, M.D., Tasic, B., Miyamichi, K., Li, L., and Luo, L. (2007). A global double-fluorescent Cre reporter mouse. Genesis 45, 593–605. 10.1002/dvg.20335.

78. Truett, G.E., Heeger, P., Mynatt, R.L., Truett, A.A., Walker, J.A., and Warman, M.L. (2000). Preparation of PCR-quality mouse genomic DNA with hot sodium hydroxide and tris (HotSHOT). Biotechniques 29, 52, 54.

79. Ewels, P.A., Peltzer, A., Fillinger, S., Patel, H., Alneberg, J., Wilm, A., Garcia, M.U., Di Tommaso, P., and Nahnsen, S. (2020). The nf-core framework for community-curated bioinformatics pipelines. Nat Biotechnol 38, 276–278. 10.1038/s41587-020-0439-x.

80. Di Tommaso, P., Chatzou, M., Floden, E.W., Barja, P.P., Palumbo, E., and Notredame, C. (2017). Nextflow enables reproducible computational workflows. Nat Biotechnol 35, 316–319. 10.1038/nbt.3820.

81. Dobin, A., Davis, C.A., Schlesinger, F., Drenkow, J., Zaleski, C., Jha, S., Batut, P., Chaisson, M., and Gingeras, T.R. (2013). STAR: ultrafast universal RNA-seq aligner. Bioinformatics 29, 15–21. 10.1093/bioinformatics/bts635.

82. Love, M.I., Huber, W., and Anders, S. (2014). Moderated estimation of fold change and dispersion for RNA-seq data with DESeq2. Genome Biol 15, 550. 10.1186/s13059-014-0550-8.

83. Robinson, J.T., Thorvaldsdottir, H., Winckler, W., Guttman, M., Lander, E.S., Getz, G., and Mesirov, J.P. (2011). Integrative genomics viewer. Nat Biotechnol 29, 24–26. 10.1038/nbt.1754.

84. Thorvaldsdottir, H., Robinson, J.T., and Mesirov, J.P. (2013). Integrative Genomics Viewer (IGV): high-performance genomics data visualization and exploration. Brief Bioinform 14, 178–192. 10.1093/bib/bbs017.

85. Elizarraras, J.M., Liao, Y., Shi, Z., Zhu, Q., Pico, A.R., and Zhang, B. (2024). WebGestalt 2024: faster gene set analysis and new support for metabolomics and multi-omics. Nucleic Acids Res 52, W415–w421. 10.1093/nar/gkae456.

