## Supplemental Data 1 for "Argonaute-2 slicing of a domesticated retrotransposon maintains proteostasis and prevents lethal skeletal muscle defects"

**Table S1**

| **$Ago2^{CD/+}$ x $Ago2^{+/–}$ (mixed C57BL/6J;N)** | | | | |
|---|---|---|---|---|
| | *$Ago2^{+/+}$* | *$Ago2^{CD/+}$* | *$Ago2^{+/–}$* | *$Ago2^{CD/–}$* |
| Observed (E15.5) | 21 | 17 | 16 | 17 |
| % observed | 30 | 24 | 23 | 24 |
| Observed (E17.5) | 6 | 9 | 14 | 10 |
| % observed | 15 | 23 | 36 | 26 |
| Observed (P0) | 37 | 30 | 27 | 6 |
| % observed | 37 | 30 | 27 | 6 |
| Observed (P21) | 52 | 42 | 43 | 0 |
| % observed | 38 | 31 | 31 | 0 |
| % expected (all ages) | 25 | 25 | 25 | 25 |

Shown are results of crossing mice on a mixed C57Bl/6J;N background. Statistical significance was determined by $X^2$ test ($P > .05$ for E15.5 and E17.5, $P = 8.9 \times 10^{-5}$ for P0, $P = 2.8 \times 10^{-10}$ for P21).

| **$Ago2^{CD/+}$ x $Ago2^{+/–}$ (50% C57BL/6J;N, 50% DBA/2J)** | | | | |
|---|---|---|---|---|
| | *$Ago2^{+/+}$* | *$Ago2^{CD/+}$* | *$Ago2^{+/–}$* | *$Ago2^{CD/–}$* |
| Observed (P0) | 14 | 17 | 19 | 5 |
| % observed | 25 | 31 | 35 | 9 |
| Observed (P21) | 35 | 28 | 37 | 0 |
| % observed | 39 | 32 | 42 | 0 |
| % expected (all ages) | 25 | 25 | 25 | 25 |

Shown are results of crossing mice on a mixed C57Bl/6J;N background with one backcross to DBA/2J. Statistical significance was determined by $X^2$ test ($P = 0.04$ for P0, $P = 1.3 \times 10^{-8}$ for P21).

| **$Ago2^{CD/+}$ x $Ago2^{+/–}$ (50% C57BL/6J;N, 50% 129X1/SvJ)** | | | | |
|---|---|---|---|---|
| | *$Ago2^{+/+}$* | *$Ago2^{CD/+}$* | *$Ago2^{+/–}$* | *$Ago2^{CD/–}$* |
| Observed (P0) | 26 | 37 | 19 | 5 |
| % observed | 30 | 43 | 22 | 6 |
| Observed (P21) | 31 | 35 | 32 | 0 |
| % observed | 32 | 36 | 33 | 0 |
| % expected (all ages) | 25 | 25 | 25 | 25 |

Shown are results of crossing mice on a mixed C57Bl/6J;N background with one backcross to 129X1/SvJ. Statistical significance was determined by $X^2$ test ($P = 1.7 \times 10^{-5}$ for P0, $P = 3.2 \times 10^{-7}$ for P21).

| *Actl6b-cre; Ago2$^{CD/+}$ x Ago2$^{fl/fl}$* | | | | |
|---|---|---|---|---|
| | *Ago2$^{+/fl}$* | *Ago2$^{fl/CD}$* | *Actl6b-Cre; Ago2$^{+/fl}$* | *Actl6b-Cre; Ago2$^{CD/fl}$* |
| Observed (P21) | 51 | 43 | 54 | 40 |
| % observed | 27 | 23 | 29 | 21 |
| % expected (all ages) | 25 | 25 | 25 | 25 |

Shown are results of crossing mice on a mixed C57Bl/6J;N background. Statistical significance was determined by $X^2$ test ($P > 0.05$).

| *Acta1-cre; Ago2$^{CD/+}$ x Ago2$^{fl/fl}$* | | | | |
|---|---|---|---|---|
| | *Ago2$^{+/fl}$* | *Ago2$^{fl/CD}$* | *Acta1-Cre; Ago2$^{+/fl}$* | *Acta1-Cre; Ago2$^{CD/fl}$* |
| Observed (P0) | 30 | 22 | 27 | 10 |
| % observed | 34 | 25 | 30 | 11 |
| Observed (P21) | 31 | 31 | 26 | 1* |
| % observed | 35 | 35 | 29 | 1 |
| % expected (all ages) | 25 | 25 | 25 | 25 |

Shown are results of crossing mice on a mixed C57Bl/6J;N background. Statistical significance was determined by $X^2$ test ($P = 0.02$ for P0, $P = 4.0 \times 10^{-6}$ for P21). Asterisk (*) denotes poor excision of the floxed allele as determined by PCR of genomic DNA from diaphragm.

| *CMV-cre; Ago2$^{CD/+}$ x Ago2$^{fl/fl}$* | | | | |
|---|---|---|---|---|
| | *Ago2$^{+/fl}$* | *Ago2$^{fl/CD}$* | *CMV-Cre; Ago2$^{+/fl}$* | *CMV-Cre; Ago2$^{CD/fl}$* |
| Observed (P0) | 15 | 6 | 15 | 3 |
| % observed | 38 | 15 | 38 | 10 |
| Observed (P21) | 20 | 7 | 24 | 2 |
| % observed | 38 | 13 | 45 | 4 |
| % expected (all ages) | 25 | 25 | 25 | 25 |

Shown are results of crossing mice on a mixed C57Bl/6J;N background. Statistical significance was determined by $X^2$ test ($P = 0.008$ for P0, $P = 1.8 \times 10^{-5}$ for P21).

| *Mlc2v-cre; Ago2$^{CD/+}$ x Ago2$^{fl/fl}$* | | | | |
|---|---|---|---|---|
| | *Ago2$^{+/fl}$* | *Ago2$^{fl/CD}$* | *Mlc2v-Cre; Ago2$^{+/fl}$* | *Mlc2v-Cre; Ago2$^{CD/fl}$* |
| Observed (P21) | 18 | 27 | 16 | 32 |
| % observed | 18 | 28 | 16 | 33 |
| % expected (all ages) | 25 | 25 | 25 | 25 |

Shown are results of crossing mice on a mixed C57Bl/6J;N background. Statistical significance was determined by $X^2$ test ($P > 0.05$).

| ***VEC-cre*** $^{+/-}$***; Ago2*** $^{CD/+}$ ***x Ago2*** $^{fl/fl}$ | | | | |
|---|---|---|---|---|
| | *Ago2* $^{+/fl}$ | *Ago2* $^{fl/CD}$ | *VEC-Cre; Ago2* $^{+/fl}$ | *VEC-Cre; Ago2* $^{CD/fl}$ |
| Observed (P21) | 23 | 34 | 19 | 30 |
| % observed | 22 | 32 | 18 | 29 |
| % expected (all ages) | 25 | 25 | 25 | 25 |

Shown are results of crossing mice on a mixed C57Bl/6J;N background. Statistical significance was determined by $X^2$ test ($P$ > 0.05 for P21).

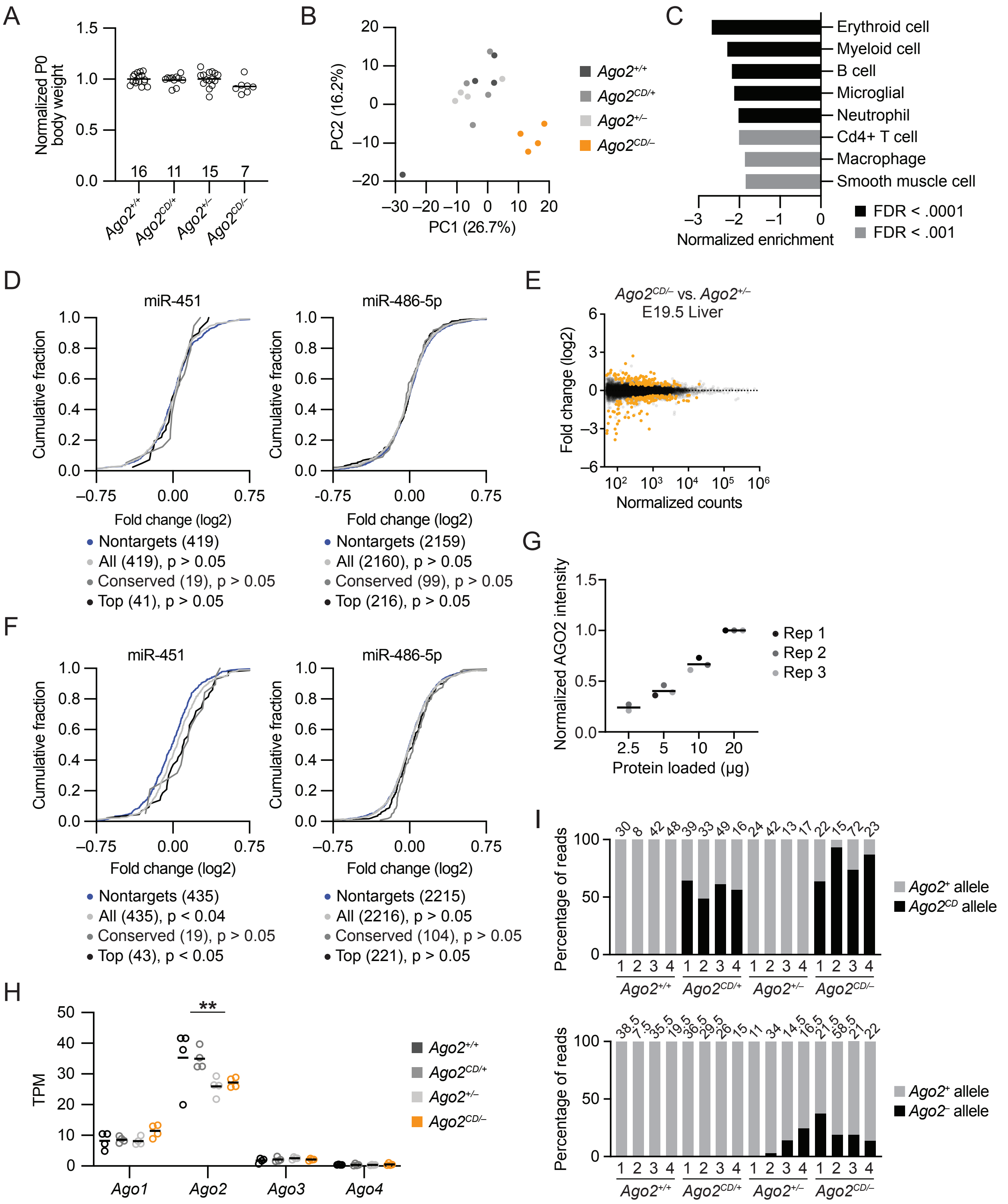
A
Normalized P0 body weight
1.5
1.0
0.5
0.0
16
11
15
7
Ago2+/+
Ago2CD/+
Ago2+/–
Ago2CD/–
B
PC2 (16.2%)
PC1 (26.7%)
20
10
0
–10
–20
–30 –20 –10 0 10 20
Ago2+/+
Ago2CD/+
Ago2+/–
Ago2CD/–
C
Erythroid cell
Myeloid cell
B cell
Microglial
Neutrophil
Cd4+ T cell
Macrophage
Smooth muscle cell
–3 –2 –1 0
Normalized enrichment
FDR < .0001
FDR < .001
D
miR-451
miR-486-5p
Cumulative fraction
Fold change (log2)
–0.75 0.00 0.75
Nontargets (419)
All (419), p > 0.05
Conserved (19), p > 0.05
Top (41), p > 0.05
Nontargets (2159)
All (2160), p > 0.05
Conserved (99), p > 0.05
Top (216), p > 0.05
E
Ago2CD/– vs. Ago2+/–
E19.5 Liver
Fold change (log2)
Normalized counts
F
miR-451
miR-486-5p
Cumulative fraction
Fold change (log2)
Nontargets (435)
All (435), p < 0.04
Conserved (19), p > 0.05
Top (43), p < 0.05
Nontargets (2215)
All (2216), p > 0.05
Conserved (104), p > 0.05
Top (221), p > 0.05
G
Normalized AGO2 intensity
Protein loaded (µg)
2.5 5 10 20
Rep 1
Rep 2
Rep 3
H
TPM
**
Ago1
Ago2
Ago3
Ago4
Ago2+/+
Ago2CD/+
Ago2+/–
Ago2CD/–
I
Percentage of reads
30 8 42 48 39 33 49 16 24 42 13 17 22 15 72 23
1 2 3 4 1 2 3 4 1 2 3 4 1 2 3 4
Ago2+/+
Ago2CD/+
Ago2+/–
Ago2CD/–
Ago2+ allele
Ago2CD allele
Percentage of reads
38.5 7.5 35.5 19.5 36.5 29.5 26 15 11 34 14.5 16.5 21.5 58.5 21 22
Ago2+ allele
Ago2– allele

**Figure S1. *Ago2*$^{CD/–}$ animals recapitulate phenotypes reported in *Ago2*$^{CD/CD}$ animals, related to Figure 1.** (A) The influence of AGO2 slicing on neonatal weight. Plotted are the normalized weights of each animal at P0 (n = 7–16 per genotype). Raw weights were divided by the average weight for each litter to account for litter-to-litter differences. No significant changes were detected ($P$ <0.05; one-way ANOVA). (B) The influence of AGO2 slicing on fetal liver gene expression. Plotted are the first two principal components (PC1, PC2) for E15.5 fetal liver based on analysis of the 500 most variable genes derived from RNA sequencing libraries. Each dot represents a unique sample/library and each genotype is represented by a different color. (C) Gene set enrichment analysis of *Ago2*$^{CD/–}$ E15.5 fetal liver. Plotted are normalized enrichment scores for significant cell-type gene sets from the Mouse Cell Atlas, as determined by WebGestalt (FDR < 0.01). (D) The influence of *Ago2*$^{CD}$ heterozygosity on expression of predicted targets of miR-451 (left) and miR-486-5p (right). Plotted are the cumulative distribution functions of mRNA fold-changes in *Ago2*$^{CD/+}$ fetal liver relative to *Ago2*$^{+/+}$ fetal liver for all predicted targets (light gray), conserved predicted targets (gray), and top 10% predicted targets (black). Otherwise, as in Figure 1E. (E) The influence of AGO2 slicing on RNA levels in the E19.5 fetal liver, as determined by RNA-seq. Otherwise, as in Figure 1D. (F) The influence of AGO2 slicing on expression of predicted targets of miR-451 (left) and miR-486-5p (right) in E19.5 fetal liver. Otherwise, as in Figure 1E. (G) Dynamic range of AGO2 protein detection in E15.5 *Ago2*$^{+/+}$ fetal liver. Plotted are AGO2 intensities, as determined by western blot of total protein from three technical replicates (separate gels), and reported relative to the intensity of AGO2 from 20 μg total protein. (H) The influence of *Ago2* heterozygosity on *Ago2* RNA levels in E15.5 fetal liver. Plotted are transcripts per million (TPM) for *Ago2* and its paralogs *Ago1*, *Ago3*, and *Ago4*. The *Ago2*$^{–}$ allele is associated with a 24% decrease in *Ago2* levels (**, $P$ < 0.01; Welch's t-test). No significant changes were detected for *Ago1*, *Ago3*, and *Ago4*. (I) Allele-specific expression of *Ago2* in E15.5 fetal liver. Plotted are the percentage of reads matching either the wild-type and *Ago2*$^{CD}$ allele (above) or the wild-type and *Ago2*$^{–}$ allele (below). For the second comparison, the *Ago2*$^{–}$ allele was quantified using splice junction reads spanning exons 9–12, whereas the wild-type allele was quantified using the average of splice junction reads spanning exons 9–10 and 11–12. Each bar represents a different animal, and the total number of reads are indicated above each bar.

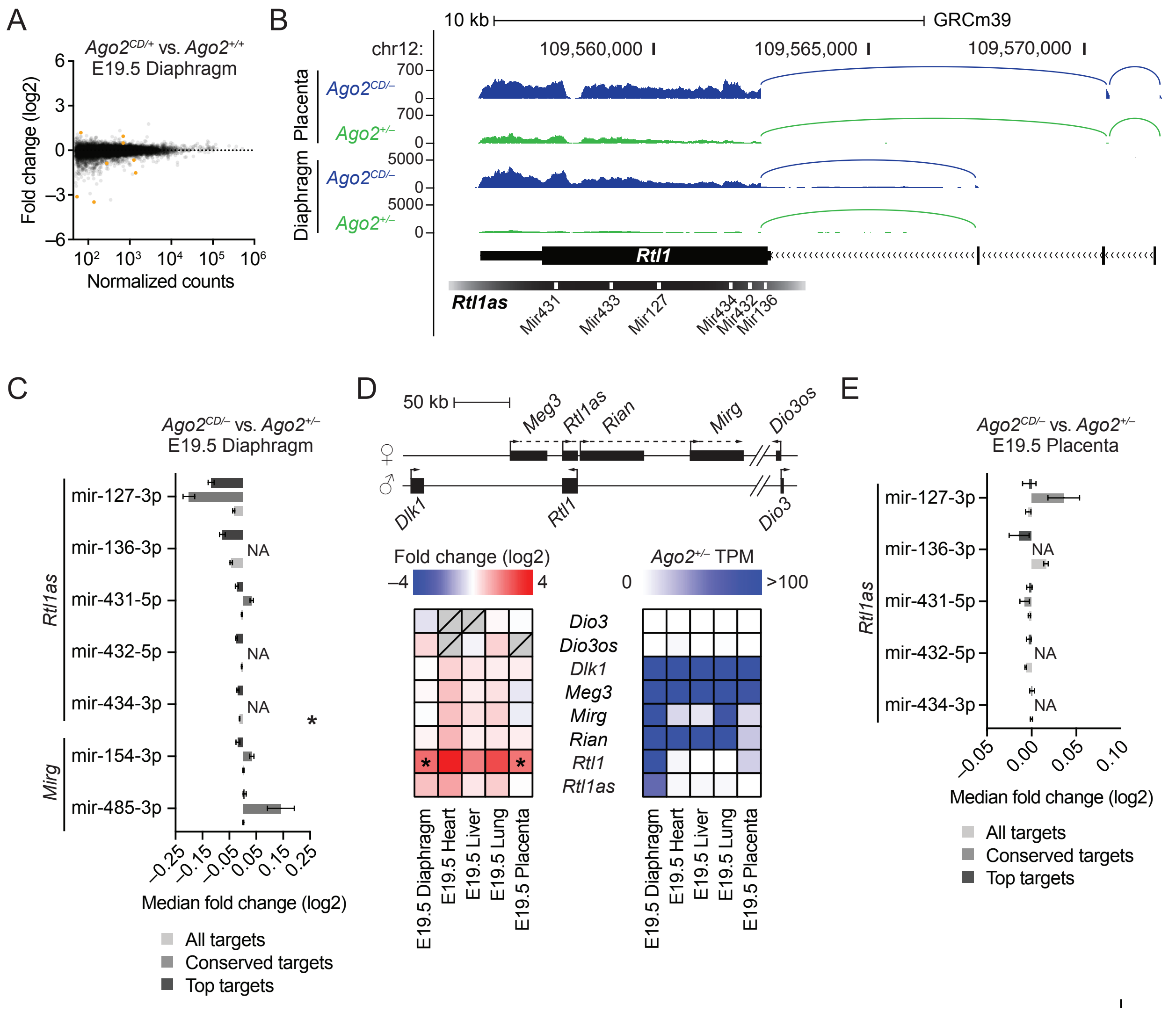

A
Ago2CD/+ vs. Ago2+/+
E19.5 Diaphragm
Fold change (log2)
Normalized counts
B
10 kb
GRCm39
chr12:
109,560,000
109,565,000
109,570,000
Placenta
Diaphragm
Ago2CD/–
Ago2+/–
Rtl1
Rtl1as
Mir431
Mir433
Mir127
Mir434
Mir432
Mir136
C
Ago2CD/– vs. Ago2+/–
E19.5 Diaphragm
mir-127-3p
mir-136-3p
mir-431-5p
mir-432-5p
mir-434-3p
mir-154-3p
mir-485-3p
Rtl1as
Mirg
NA
Median fold change (log2)
All targets
Conserved targets
Top targets
D
50 kb
Meg3
Rtl1as
Rian
Mirg
Dio3os
Dlk1
Rtl1
Dio3
Fold change (log2)
Ago2+/– TPM
>100
Dio3
Dio3os
Dlk1
Meg3
Mirg
Rian
Rtl1
Rtl1as
E19.5 Diaphragm
E19.5 Heart
E19.5 Liver
E19.5 Lung
E19.5 Placenta
E
Ago2CD/– vs. Ago2+/–
E19.5 Placenta
mir-127-3p
mir-136-3p
mir-431-5p
mir-432-5p
mir-434-3p
Rtl1as
NA
Median fold change (log2)
All targets
Conserved targets
Top targets

**Figure S2: AGO2 endonuclease activity affects gene expression and substrates in multiple tissues, related to Figure 2.** (A) The influence of *Ago2*$^{CD}$ heterozygosity on RNA levels in E19.5 diaphragm (n = 3 per genotype). Otherwise, as in Figure 2A. (B) Organization and expression of the *Rtl1* locus on chromosome 12 (GRCm39). Shown are gene models for *Rtl1* and *Rtl1as* (black boxes, exons; >>>, introns; white boxes, microRNA hairpins); the precise coordinates for *Rtl1as* have not been established. Shown above, the gene model are RNA-seq tracks from E19.5 *Ago2*$^{CD/–}$ and *Ago2*$^{+/–}$ placenta and diaphragm (y-axis, reads) derived from libraries of similar sequencing depth. Splice junctions were manually annotated based on junction-spanning reads. Distinct *Rtl1* transcript isoforms were detected in diaphragm and placenta. (C) Influence of AGO2 slicing on expression of predicted targets of *Rtl1as*-derived and *Mirg*-derived miRNAs in E19.5 diaphragm. For microRNAs that are poorly conserved, analysis of conserved targets was omitted and denoted NA. Otherwise, as in Figure 2B. (D) Organization and expression of genes in the imprinted *Dlk1-Dio3* locus (black boxes, genes; hashmarks, ~500 kb interval between *Mirg* and *Dio3*/*Dio3os*). Below the schematic, heatmaps indicate fold-changes in *Ago2*$^{CD/–}$ tissue relative to *Ago2*$^{+/–}$ tissue (left) and mean transcripts per million (TPM) in *Ago2*$^{+/–}$ tissue (right) for the indicated genes. Otherwise, as in Figure 2B. (E) Influence of AGO2 slicing on expression of predicted targets of *Rtl1as*-derived miRNAs in E19.5 placenta. Otherwise, as in Figure 2C.

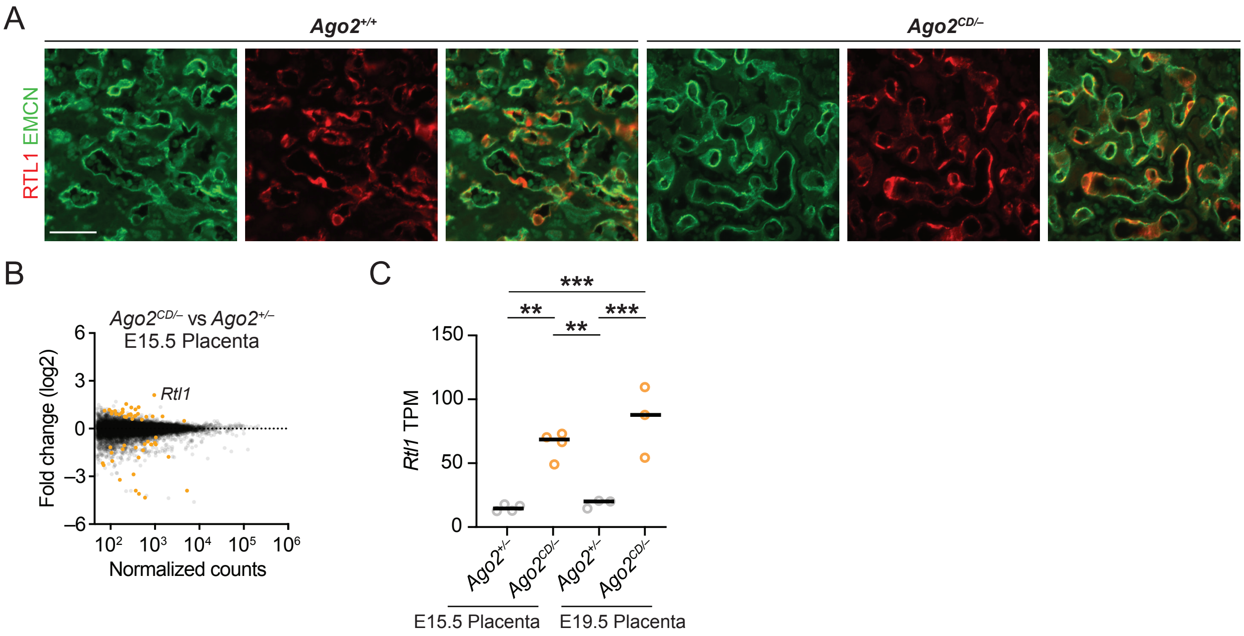

A
Ago2+/+
Ago2CD/–
RTL1 EMCN
B
Ago2CD/– vs Ago2+/–
E15.5 Placenta
Fold change (log2)
Rtl1
Normalized counts
C
Rtl1 TPM
Ago2+/–
Ago2CD/–
Ago2+/–
Ago2CD/–
E15.5 Placenta
E19.5 Placenta

**Figure S3: Increased *Rtl1* in *Ago2*$^{CD/–}$ placenta does not cause placentomegaly or vascular defects, related to Figure 3.** (A) Influence of AGO2 slicing on RTL1 localization, as determined by immunofluorescence for Endomucin (EMCN, green) and RTL1 (red). Shown are representative images of fetal vasculature from E19.5 *Ago2*$^{+/+}$ and *Ago2*$^{CD/–}$ placenta; scale bar, 20 microns. (B) The influence of AGO2 slicing on RNA levels in the E15.5 fetal placenta (n = 4 per genotype). Otherwise, as in Figure 1D. (C) *Rtl1* TPMs in E15.5 and E19.5 placenta (**, $P < 0.01$; ***, $P < 0.001$; one-way ANOVA with Tukey's multiple comparisons test).

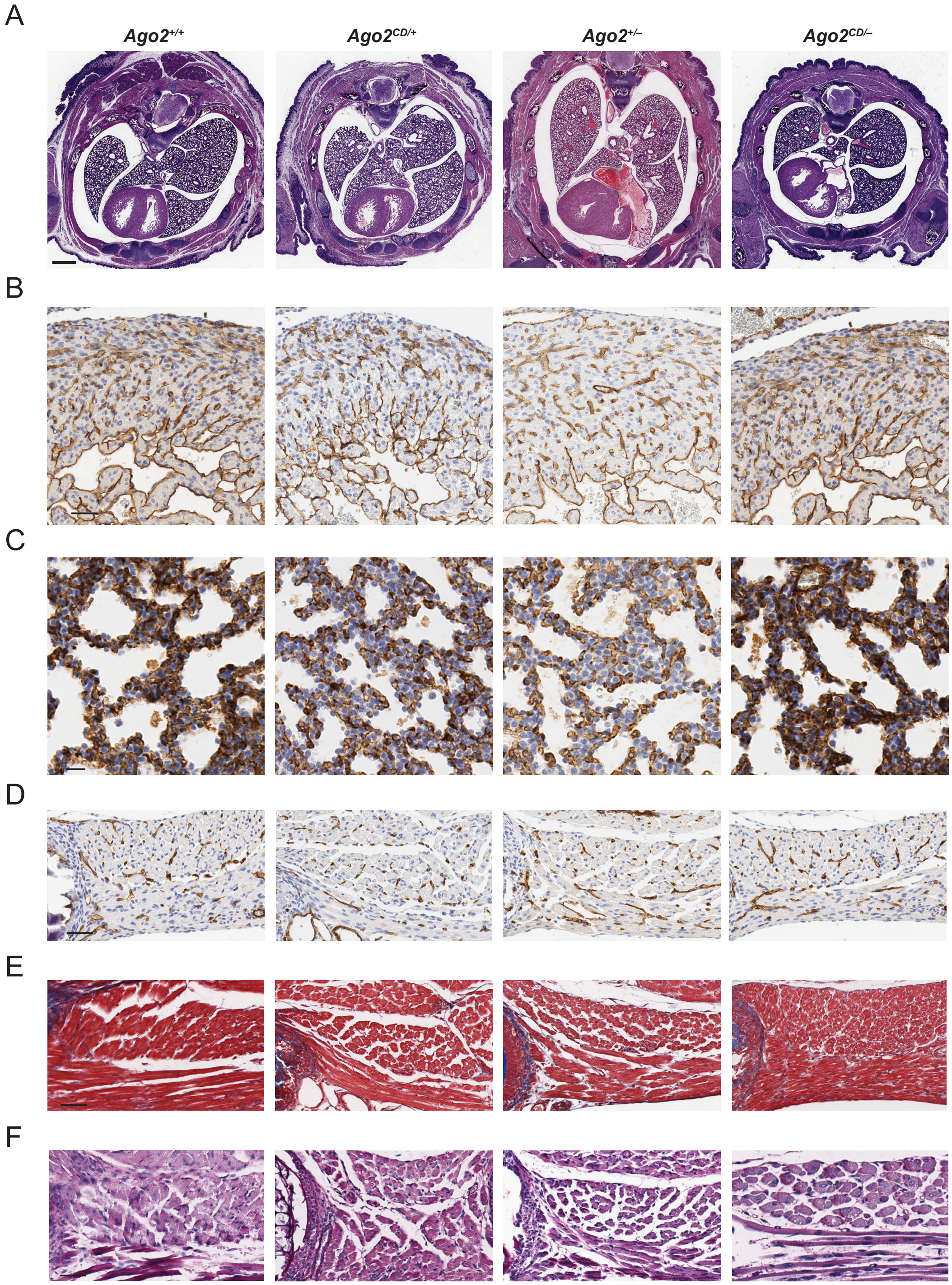
A
Ago2+/+
Ago2CD/+
Ago2+/–
Ago2CD/–
B
C
D
E
F

**Figure S4: *Ago2*$^{CD/–}$ skeletal muscle fibers are larger and contain more central nuclei, related to Figure 4.** (A) The influence of AGO2 slicing on embryonic development, as determined by hematoxylin and eosin staining. Shown are representative images of transverse sections from E19.5 mouse torso; scale bar, 800 microns. (B–D) The influence of AGO2 slicing on embryonic vasculature, as determined by immunohistochemistry for PECAM-1 (brown). Shown are representative images of E19.5 right ventricle (B), lung (C), and intercostal muscle (D); scale bar, 50 microns in panels B and D, and 20 microns in panel C. (E) The influence of AGO2 slicing on collagen accumulation, as determined by Masson's trichrome staining; scale bar, 50 microns. (F) The influence of AGO2 slicing on glycogen accumulation, as determined by Periodic Acid-Schiff staining; scale bar, 50 microns.

A

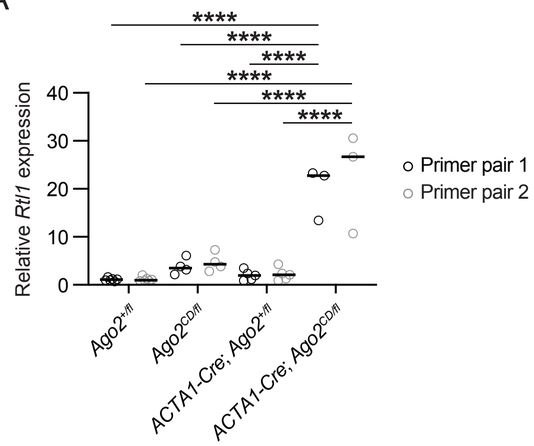

****
****
****
****
****
****
Relative Rtl1 expression
40
30
20
10
0
Ago2+/fl
Ago2CD/fl
ACTA1-Cre; Ago2+/fl
ACTA1-Cre; Ago2CD/fl
Primer pair 1
Primer pair 2

**Figure S5: Loss of AGO2 endonuclease activity in skeletal muscle causes muscle defects and perinatal lethality, related to Figure 5.** (A) Relative abundance of *Rtl1* in E19.5 diaphragm, as determined by RT-qPCR using two pairs of *Rtl1* primers specific to the skeletal muscle-enriched *Rtl1* transcript isoform. Levels of *Rtl1* were normalized to the geometric mean of *Actb*, *Gapdh*, and *Acta1* and plotted relative to mean $Ago2^{+/fl}$ expression for each primer pair (n = 3–6 per genotype; ****, $P < 0.0001$; one-way ANOVA with Tukey's multiple comparisons test).

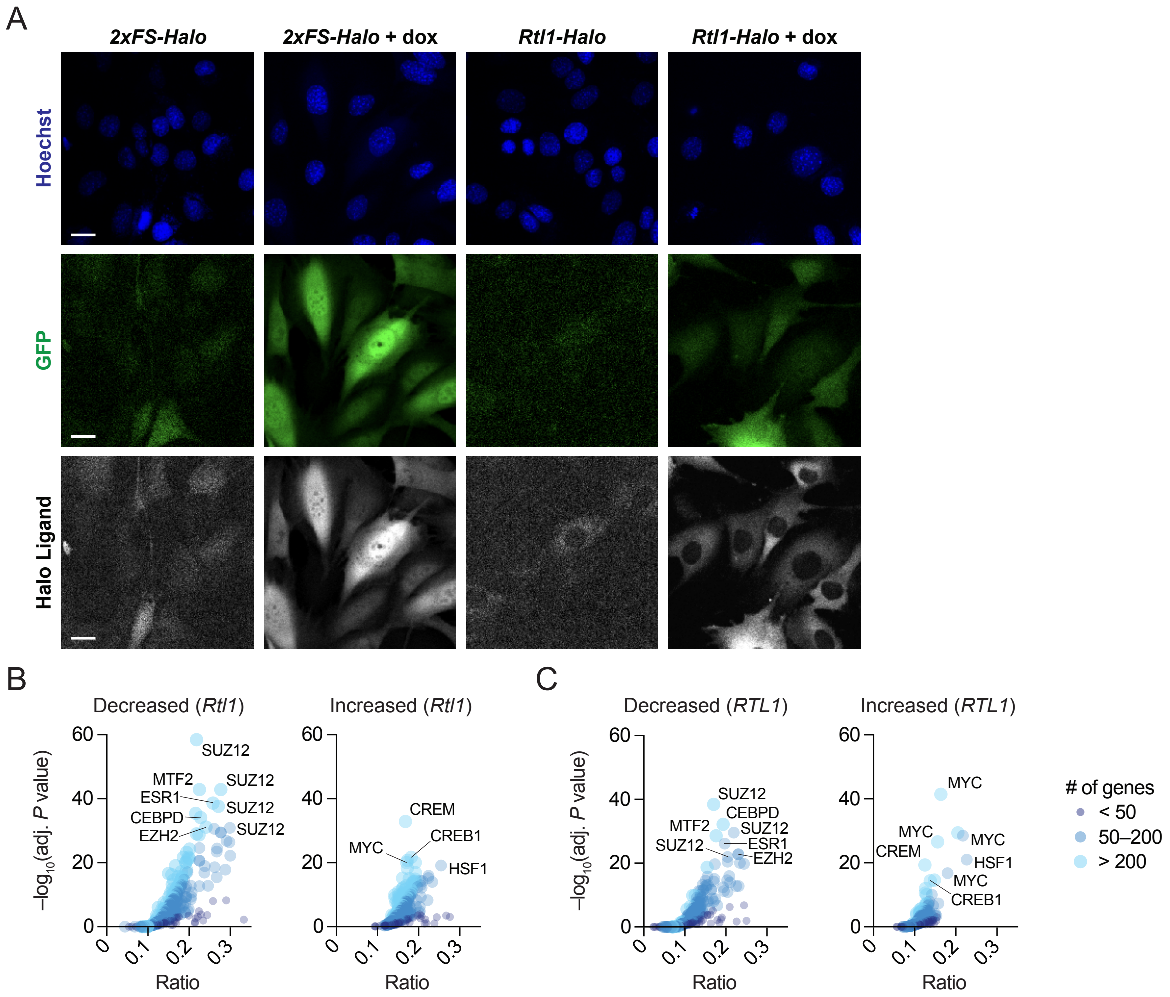
A
2xFS-Halo
2xFS-Halo + dox
Rtl1-Halo
Rtl1-Halo + dox
Hoechst
GFP
Halo Ligand
B
Decreased (Rtl1)
Increased (Rtl1)
C
Decreased (RTL1)
Increased (RTL1)
−log10(adj. P value)
Ratio
SUZ12
MTF2
ESR1
CEBPD
EZH2
CREM
MYC
CREB1
HSF1
### of genes
< 50
50–200
> 200

**Figure S6: RTL1 expression induces a heat shock response in muscle cells, related to Figure 6.** (A) Subcellular localization of 2xFS-Halo or mouse RTL1-Halo in C2C12 myoblasts, as determined by live-cell confocal microscopy. Shown are representative images of untreated and doxycycline-treated *2xFS-Halo*- and *Rtl1-Halo*-expressing cell lines after staining with Hoechst (nuclei) and JF-646 Halo ligand; each cell line also expresses GFP from the same expression cassette via a T2a peptide; scale bar, 20 microns. As detailed in the methods section, the detection parameters were adjusted for the doxycycline-treated *2xFS-Halo*-expressing cell line to avoid saturating the image. (B–C) Overrepresentation of transcription factor targets in differentially expressed gene sets from *Rtl1/RTL1*-expressing C2C12 myoblasts, as determined by Enrichr. Plotted are the ratio of transcription factor targets present among DEGs that decrease (left) or increase (right) with *Rtl1* (B) or *RTL1* (C) induction versus the adjusted *P*-value. Each circle represents a set of high-confidence targets for one transcription factor, based on the 2022 ChIP-X Enrichment Analysis database, with the size of the circle reflecting the number of targets that overlap with the set of increased or decreased DEGs.
